# Spatially precise neuron formation via hydrogel mediated modulation of the host astrocyte response

**DOI:** 10.64898/2025.12.20.695501

**Authors:** Negar Mahmoudi, Alan Harvey, Niamh Moriarty, Morteza Mahmoudi, Wei Tong, Toon Goris, Nathan Reynolds, Noorya Y. Ahmed, Nathalie Dehorter, Leszek Lisowski, Clare L. Parish, Richard J. Williams, David R. Nisbet

## Abstract

The regenerative capacity of the central nervous system after injury or disease is limited. Advancements in genetic and epigenetic reprogramming of non-neuronal cells into induced neurons presents a promising strategy for neuronal replacement and circuit reconstruction yet is hindered by suboptimal reprogramming efficacy. Here, we have achieved high astrocyte-to-neuron reprogramming efficacy through the ectopic expression of a SOX2 transcription factor using adeno-associated viral vectors. We subsequently designed a hybrid composite biomaterial to act as an implantable reprogramming workshop that perform sequential operations. Once injected, this bespoke system forms a tissue mimetic hydrogel encouraging the ingress of astrocytes to act as our raw material. We demonstrate that the system localized and enhanced the spatially confined delivery of reprogramming factors, entrapping astrocytes to then pass through an intermediate neuroblast state to robustly yield mature neurons. Sustained delivery of valproic acid within the system further promoted neuronal maturation. Overall, using design rules informed by requirements of the reprogramming strategy, we have designed a smart delivery system optimized for both the target tissue and reprogramming factors in a minimally invasive manner to enhance neural repair.

**Graphical Abstract:** 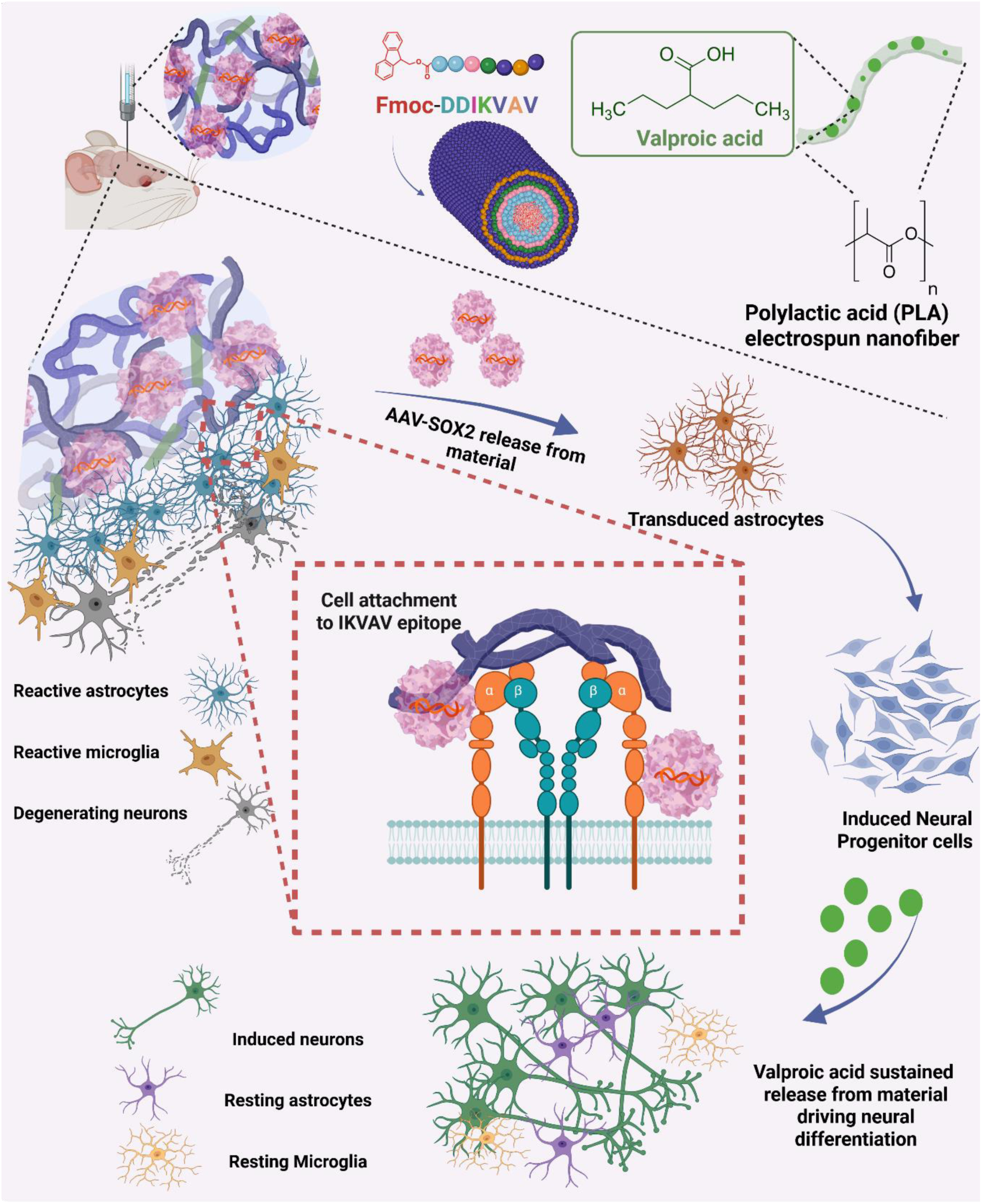

## Introduction

Neural injury, as a consequence of insult or degenerative processes can have a plethora of consequences on motor, sensory and cognitive function with a significant impact on individuals, carers and the broader socioeconomic landscape^1^. Unfortunately, with limited capacity for regeneration, outcomes are often bleak especially for sufferers of neurodegenerative diseases, stroke or other forms of brain injury. Necessary for recovery is the replacement of lost neural circuitry, and whilst neural transplantation is currently being explored, an alternative approach in recent years has been the ability to reprogram host cells into new replacement neurons.

In response to neural injuries and degeneration, astrocytes become reactive, undergoing significant morphological^2^, transcriptional, and functional changes^3^. Studies show that up to 50% of astrocytes re-enter the cell cycle and proliferate within three days to a week post-injury, forming a protective barrier known as the glial scar^4^. The scar provides several important beneficial functions such as confining the lesion and preventing its spread, promoting repair of the blood-brain barrier (BBB), and limiting neuronal loss in ischemic stroke and neurotrauma^5^. However, the presence of mature glial scars that persist for a long period act as a barrier to axonal regrowth and extension beyond the site of the initial injury resulting in dystrophic end balls. These scars also negatively affect the successful integration of new neural grafts into host tissue^6^. Therefore, methods acting as multifaceted tools to limit gliosis and inflammatory responses, after they have exerted their protective role may allow the re-establishment of lost neural circuitry and aid in some level of functional recovery^7–12^. Given astrocytes’ abundance throughout the brain, unlike the limited pool of neural progenitors (which are largely confined to neurogenic zones)^13–15^, reprogramming of reactive astrocytes offers a promising strategy for generating new neurons^16^. This glia can generate new neurons if genetically reprogrammed by master regulator/pioneer transcription factors (TFs)^9–11,17–21^.

Recent advances have shown that somatic cells can be reprogrammed into pluripotent states^22,23^ by introducing a specific set of TFs including OCT-3/4, SOX2, KLF4, and C-MYC^24^. However, this approach has led to concerns about the potential for inducing tumorigenesis *in vivo*^25,26^. As an alternative, it has been shown that SOX2 can reprogram astrocytes into induced progenitor neural stem cells (iPNSCs)/ or neural stem cells (NSCs)^21,27^. This approach mitigates some of the risks associated with reverting cells into induced pluripotent stem cells (iPSCs) whereby teratoma formation and incomplete differentiation can occur^28,29^. This lineage-restricted stem cell reprogramming complements the iPSC technology as no histological signs of tumorigenesis have been observed in brains and/or spinal cords with ectopic SOX2 expression up to one-year post lentiviral vector injection^21^. *In vivo*, this ectopic expression initiates a progressive reprogramming process, generating neuronal progenitor cells/neuroblast cells and ultimately mature neurons, positive for NeuN marker, when supplemented with brain-derived neurotrophic factor (BDNF) or valproic acid (VPA, a histone deacetylase (HDAC) inhibitor)^30^, highlighting the crucial role of the microenvironment in influencing cell differentiation and reprogramming in adult CNS^31^.

For these reasons, biomaterials that can engage with the surrounding tissue and support mature lineage commitment, present a valuable and readily exploitable resource to enhance the reprogramming process^32^. Examining the interplay between cells and their microenvironments is essential for developing next-generation biomaterials that guide cell reprogramming effectively. Three-dimensional (3D) biomaterial-based microenvironments offer robust design options that allow control over the biophysical (including, but not limited to, topography, stiffness and elasticity, and mechanical/extracellular forces) and biochemical (including extracellular matrix (ECM) components, small molecules, soluble factors, growth factors, and other signaling molecules) cues that are known to modulate cell fate and cell reprogramming^33–35^. Stiffness is particularly important, as it influences stem cell differentiation^36–40^; stiffer substrates promote osteogenic differentiation^41,42^, while softer matrices favour neuronal or adipocyte differentiation^41^. Stiffness promotes cell tension (changes in actin force) and thus activation of biological and chemical pathways^43^ (e.g., chromatin conformation) that lead to reprogramming into appropriate phenotype^44^. In this study, a N-fluorenylmethyloxycarbonyl self-assembling peptide (Fmoc-SAPs) hydrogel was selected as the underlying matrix, as it can mimic brain ECM stiffness and promote cell adhesion while concomitantly presenting the IKVAV (isoleucine-lysine-valine-alanine-valine) peptide sequence mimicking laminin. This matrix was chosen to be delivered via direct injection to promote cell interaction, attenuate glial scar formation, and support host astrocytes infiltration^18,45^. The non-covalent structure of these hydrogels was used to encapsulate adeno-associated viral vectors (AAV). Then, VPA loaded electrospun nanofibers were immobilized in the SAP hydrogel to fabricate a hybrid composite biomaterial to provide sustained delivery for VPA to obviate the need for daily intraperitoneal (i.p.) injection to direct induced neurons (iNs) towards a specific differentiation pathway.

We examined the synergistic effect of multiple approaches including gene delivery, biomaterials, and small molecules to generate iNs and further enhance their differentiation and maximize the regenerative potential and reprogramming efficiency. This multifaceted approach targets various aspects: i) Fmoc-SAP hydrogel provides physical and chemical cues to support cell differentiation; ii) localized delivery of a gene therapy resulted in the ectopic expression of SOX2 which activates regenerative genes, reducing glial scar formation and promoting neuronal regeneration; and iii) the hybrid composite biomaterial offers spatiotemporal release of VPA, reducing secondary injury events and driving iPNSC differentiation into mature neurons.

## Results and Discussion

### AAV-SOX2 Dosage Optimization *In vitro*

First, to ensure an effective reprogramming methodology we employed an AAV delivery system to target astrocytes for cell fate reprogramming. To avoid off-targeting transduction, we utilized a cell-specific serotype (DJ) and astrocyte targeted promoter, which enhanced the expression of the transgene specifically in target cells^18,46,47^. Thus, a truncated version of hGFAP promoter, gfaABC1D, was constructed to selectively drive the expression of the AAVs in mature astroglial cells *in vivo/in vitro*^48,49^. Furthermore, the AAV was tagged with mCherry to enable tracking of transduced cells. The ability of the constructed AAV to preferentially transduce astrocytes, was confirmed by co-culturing astrocytic and non-astrocytic cells (microglia and neurons) and treating them with DJ gfaABC1D-mCherry vector (AAV-mCherry). The co-localization of GFAP (astrocyte marker) and mCherry and absence of transduction of other cell types confirmed the successful selective transduction of cultured astrocytic cells (Supporting Information (SI) **Figure 1SA)**.

**Figure 1.**
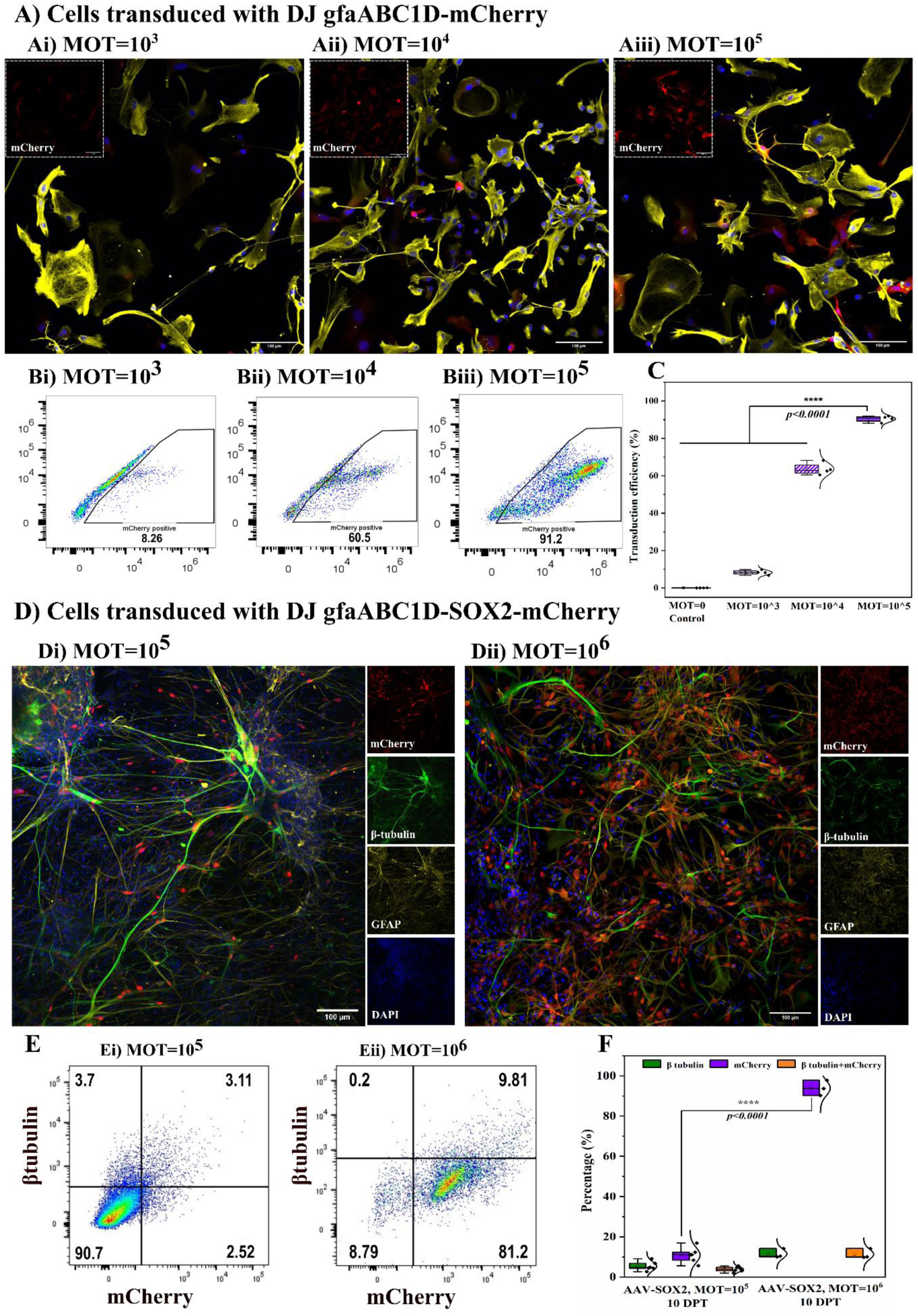
Transduction efficiency of DJ gfaABC1D-mCherry and DJ gfaABC1D-SOX2-mCherry vectors. Primary astrocytes were transduced with DJ gfaABC1D-mCherry at **Ai)** MOT=10^3^, **Aii)** MOT=10^4^ and **Aiii)** MOT=10^5^, fixed and stained with transduction efficiency marker mCherry (red), astrocytic protein GFAP (yellow), and DAPI (blue). **B)** Representative data of flow cytometry plots for DJ gfaABC1D-mCherry transduced astrocyte with different MOTs. **C)** Quantification data for mCherry+ cells. **D)** Primary astrocytes were transduced with DJ gfaABC1D-SOX2-mCherry at **Di)** MOT=10^5^ and **Dii)** MOT=10^6^, fixed and stained with mCherry (red), neuronal marker β3-tubulin (green), GFAP (yellow), and DAPI (blue) at 10 DPT. **E)** Representative data of flow cytometry plots for DJ gfaABC1D-SOX2-mCherry transduced astrocyte with MOT=10^5^ and MOT=10^6^. **F)** Quantification data for mCherry+, β-tubulin+, and both mCherry+ and β3-tubulin+ cells. Data represents Mean ± SEM, **** p < 0.0001. Scale bar=100 µm.

The constructed DJ gfaABC1D-SOX2-mCherry (AAV-SOX2-mCherry) and AAV-mCherry was tested *in vitro* on cultured murine primary astrocytes. First, we examined the purity of cultured astrocytes by staining the cells against GFAP, Iba-1 (microglia marker), and β3-tubulin (neuronal marker). The results indicated a high purity of astrocytes (94.6 ± 2.5%) in culture (**Figure 1SB-D)**. Then to optimise the transduction efficiency of the AAV-mCherry, the primary astrocytes were transduced with different multiplicities of transduction (MOT), MOT=10^3^, MOT=10^4^, and MOT=10^5^. As revealed through immunocytochemistry (**Figure 1A**) and flow cytometry analysis (**Figure 1B-C**) transduction efficiencies were 8.3 ± 1.2%, 63.6 ± 3.3%, and 90.5 ± 1.7%, for MOT=10^3^, 10^4^, 10^5^, respectively confirming the highest transduction efficiency achieved in MOT=10^5^. Therefore, MOT=10^5^ was used for AAV-mCherry transduction for subsequent experiments.

To examine the transduction and reprogramming efficiency of the AAV-SOX2-mCherry construct, the primary astrocytes were transduced with MOT=10^5^ and MOT=10^6^ of AAV-SOX2-mCherry. At three days post-transduction (DPT), the transduced astrocytes began to divide and adopt pseudoepithelial morphology. Then, to determine the transduction and reprogramming efficiency, the cells were fixed and stained at 10 DPT. Strong mCherry expression confirmed the successful transduction of astrocytes with AAV-SOX2 **(Figure 1D)**. Flow cytometry analysis was used to quantify the number of transduced and converted cells **(Figure 1E)**. Some cells were immunopositive for both β3-tubulin and mCherry (**Figure 1D and E**), which may suggest they were in a transitional stage between astrocytes and neurons. As illustrated in flow cytometry plots **(Figure 1E, F)**, the percentage of cells expressing mCherry transduction efficiency marker in MOT=10^6^ (93.9 ± 3.7% mCherry+) was significantly higher than MOT=10^5^ (10.9 ± 3.7% mCherry+). Therefore, MOT=10^6^ was selected for subsequent *in vitro* testing.

### SOX2 Transduced Astrocytes Pass Through Proliferative State

Reprogramming cells towards multipotency involves expression of master stem cell factors, initiating a cascade of molecular events that generate various intermediate cells. These regenerated progenitor cells are capable of self-renewal and can further differentiate into neurons and all macroglial cells *in situ*, effectively filling tissue cavities and/or replenishing neurons lost due to disease or injury^50^. Here, we confirmed that most SOX2-transduced cells transitioned into neural progenitor/stem cells and underwent self-renewal. The terms NSCs and NPSCs are often used interchangeably due to their limited self-renewal properties and range of progeny. The formation of iNs was validated by: i) immunostaining of nestin, an intermediate filament protein highly enriched in neural stem/ progenitor cells and lack of OCT 3/4 immunoreactivity; ii) immunostaining of Ki67, to visualize the dividing cells, iii) morphological appearance using IncuCyte live cell imaging; and iv) cell cycle analysis via flow cytometry. These methods collectively confirmed the successful reprogramming of astrocytes into neuronal progenitors.

To validate conversion to neural progenitors, astrocytes were transduced with either AAV-SOX2-mCherry or AAV-mCherry as control, fixed and immunostained for nestin. At 5 DPT, 65.7 ± 7.5% of AAV-SOX2 transduced cells were positive for nestin, while in AAV-mCherry transduced cells only 3.4 ± 1.2% were positive **(Figure 2A-C)**. This data suggests that AAV-SOX2 enables astrocytes to dedifferentiate toward iNs. The AAV-SOX2-mCherry transduced cells were also stained against Oct3/4 (pluripotency marker) and compared to the pluripotent stem cells (PSCs) as positive control to confirm the formation of induced neural progenitor (**Figure 2D,E**). The results confirm the successful reprogramming of reactive astrocytes to neural lineage-restricted neural progenitor cells (NPCs) and not PSCs. Moreover, at 10 DPT the transduced cells were stained for ki67 (to visualize proliferative cells). As expected, cells transduced with AAV-SOX2-mCherry exhibited significant cell proliferation (**Figure 2G**, Ki67+ cells, Cyan) compared to AAV-mCherry transduced cells (**Figure 2F**).

**Figure 2.**
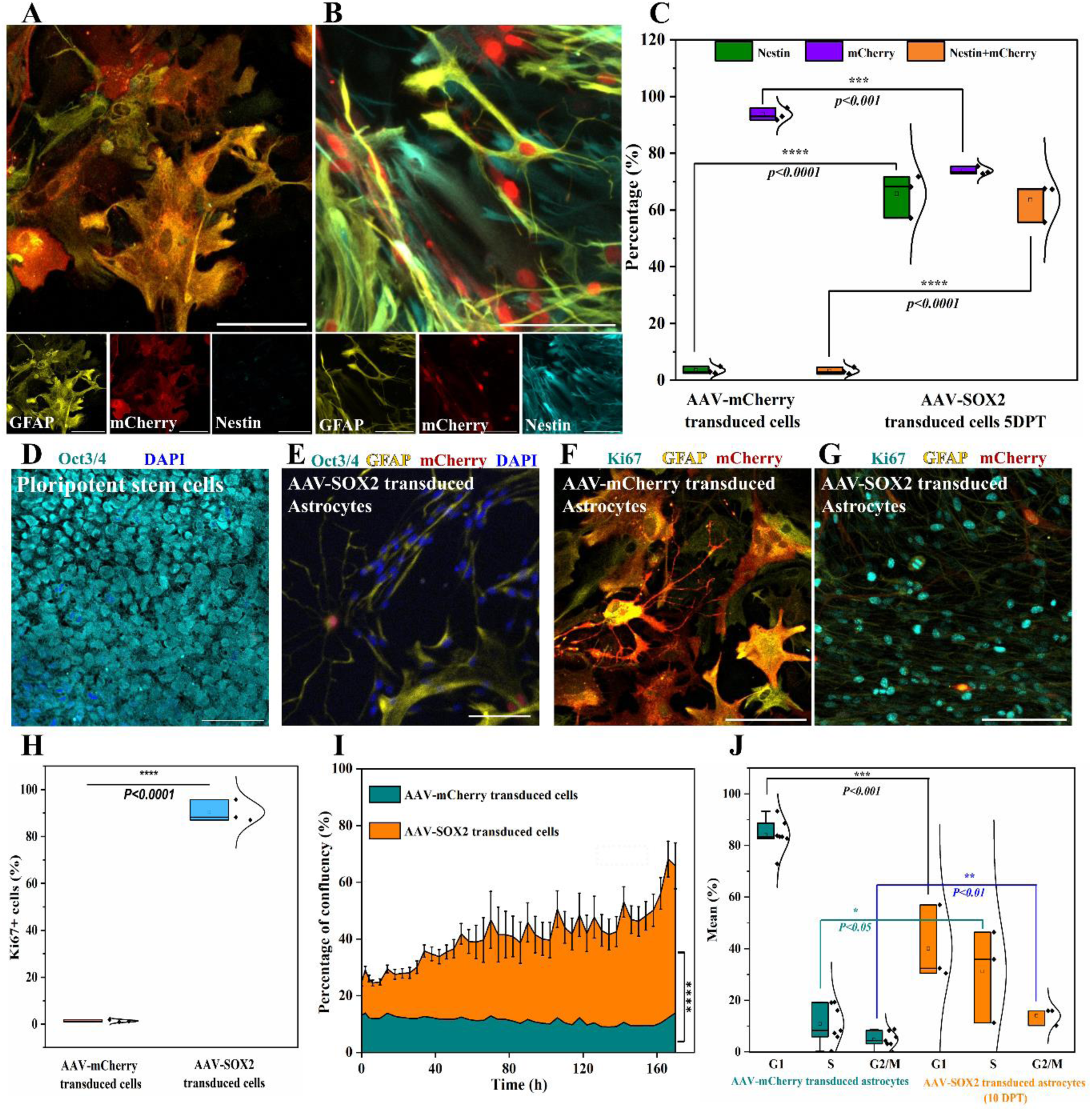
Fluorescent/live microscopy and cell cycle analysis for AAV-SOX2-mCherry and AAV-mCherry transduced cells. A representative image of astrocytes transduced with **A, F)** AAV-mCherry and **B, E, G)** AAV-SOX2 and stained with nestin/OCT3-4/Ki67 marker (cyan), mCherry (red), GFAP (yellow), and DAPI (blue). **C)** Quantification data showing the population of nestin+, mCherry+, and double positive cells (both nestin+ and mCherry+) in culture. **H)** Quantification data showing the population of ki67 positive cells, proliferation marker, in AAV-mCherry and AAV-SOX2-mCherry transduced cells. **I)** The percentage of confluence for AAV-SOX2-mCherry and AAV-mCherry transduced cells at any given time point acquired using IncuCyte live imaging. **J)** Cell cycle analysis on AAV-SOX2-mCherry and AAV-mCherry transduced astrocytes showing significant increase in S phase and G2/M phase of SOX2 transduced cells indicating the presence proliferating cells. Data represents Mean ± SEM, *\* p < 0.05, ** p < 0.01, *** p < 0.001, **** p < 0.0001*. N=3-5 independent cultures/group. Scale bar=100 µm.

After initial transduction of astrocytes with AAV-SOX2-mCherry or AAV-mCherry, cells were monitored for 7 days using live microscope IncuCyte, capturing images every 4 hours. The percentage of confluent cells at any given time point was calculated **(Figure 2I)**. As expected, the cells that were transduced with AAV-SOX2-mCherry (**Movie SIA)** exhibited significant greater cell proliferation compared with AAV-mCherry transduced control cultures **(Movie SIB)**. This observation suggests that the cell conversion using SOX2 is not a linear process with the potential of one astrocyte being reprogrammed and giving rise to multiple NPSCs. To further characterize these cells molecularly, cell cycle analysis was performed. Transducing astrocytes with episomal AAV-SOX2-mCherry during the G1 phase and transitioning them through the S phase can generate a pool of iNs. To investigate this, astrocytes were transduced with either AAV-SOX2-mCherry or AAV-mCherry and their cell cycle stages were examined at 10 DPT using flow cytometry. The process of cell proliferation and division was confirmed by observing cells progressing through the S phase, which is indicative of DNA synthesis and replication, and the G2/M phase, which indicates DNA duplication and preparation of cell division. As revealed in **Figure 2J** AAV-SOX2-mCherry transduced cells re-entered the S phase, indicating an up-regulation of cell cycle components associated with their proliferation and division. Significant differences were noted in the G0-G1, S, and G2/M phases between AAV-SOX2-mCherry and AAV-mCherry transduced cells. Collectively, data from nestin marker, IncuCyte live imaging, and cell cycle analysis indicate that the desired cells (iNs) were successfully induced.

### *In vitro* Transduction of Primary Astrocytes with SOX2 Leads to Astrocyte to Neuron Conversion

A crucial objective in the reprogramming of terminally differentiated astrocytes is to produce sufficient numbers of functional neurons. At 10 DPT, as revealed by microscopic images and flow cytometry data **(Figure 1D-F)**, the percentage of cells expressing β3-tubulin was 11.4 ± 2.3%, considered a low reprogramming efficiency (<20%)^51^. Therefore, a longer period was required to achieve higher reprogramming efficiencies.

During dedifferentiation, the ongoing presence of BDNF or VPA is of paramount importance to further drive the maturation of newly generated cells^52^. We initially examined the effect of using BDNF or VPA *in vitro* on primary astrocytes by supplementing these two candidate factors into the differentiation medium. Cells were maintained for 28 DPT with medium changed every 3-4 days. Control samples were divided into four different groups i) astrocytes transduced with empty vector and treated with VPA, ii) astrocytes treated with VPA (without any vector transduction), iii) astrocytes transduced with empty vector and treated with BDNF, and vi) astrocytes treated with BDNF (without any vector transduction) for 28 days. Controls and AAV-SOX2-mCherry transduced cells were fixed and stained for mCherry+ (to visualize AAV), GFAP+ (to visualize astrocytes), and β3-tubulin+/ MAP2+/ NeuN+ (to visualize induced neurons).

As seen in control samples **(Figure 3A)**, whether transduced or not, BDNF or VPA treatment had no noticeable impact on the morphology of cultured astrocytes **(Figure 3B)**. Moreover, the quantification data **(Figure 3G-L)** acquired using flow cytometry did not reveal any significant differences between groups (**Figure 3D, E**). These data suggest that either VPA or BDNF alone are not sufficient to induce astrocyte conversion in culture. In comparison, in AAV-SOX2-mCherry transduced cells at 28 DPT, many cells exhibited neuronal-like phenotypes (small round soma with at least 2 neuritic processes) after the addition of BDNF or VPA treatment **(Figure 3C)**, with significantly greater efficiency confirmed by flow cytometry analysis (cells positive for β3-tubulin marker, AAV-SOX2-mCherry + BDNF: 68.7 ± 7% and AAV-SOX2-mCherry + VPA: 65.2 ± 5.8%) (**Figure 3G-L**). Cells were also MAP2+ with 68.6 ± 1.3% for BDNF treated and 68.4 ± 0.6% for VPA treated. Moreover, 62.4 ± 2.1% and 61.3 ± 2.6% of cells were NeuN+, in BDNF and VPA supplemented medium, respectively. Interestingly, no significant statistical difference was observed between efficiency of reprogramming between BDNF and VPA treated cells (one-way *ANOVA,* p<0.05).

**Figure 3.**
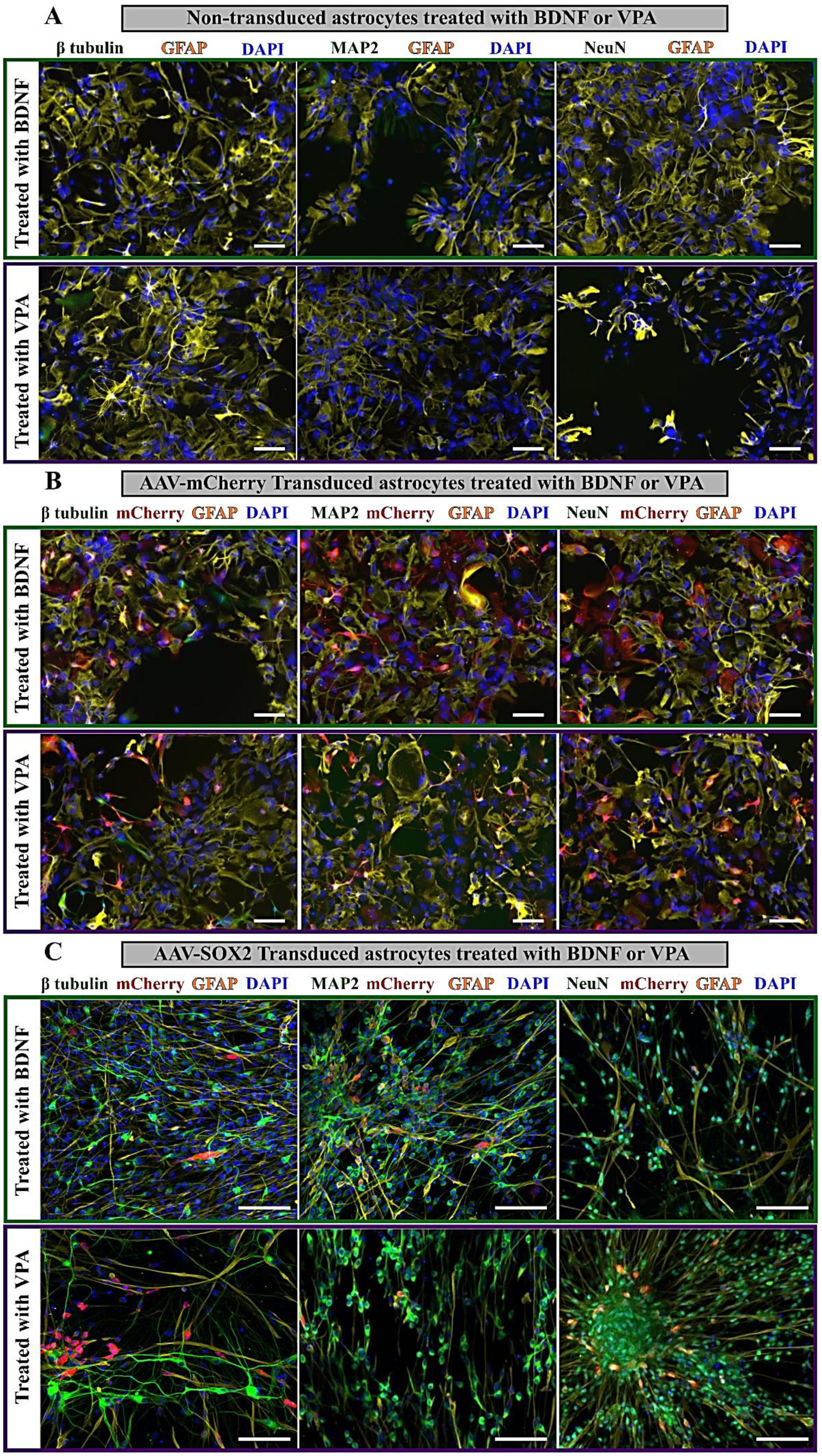
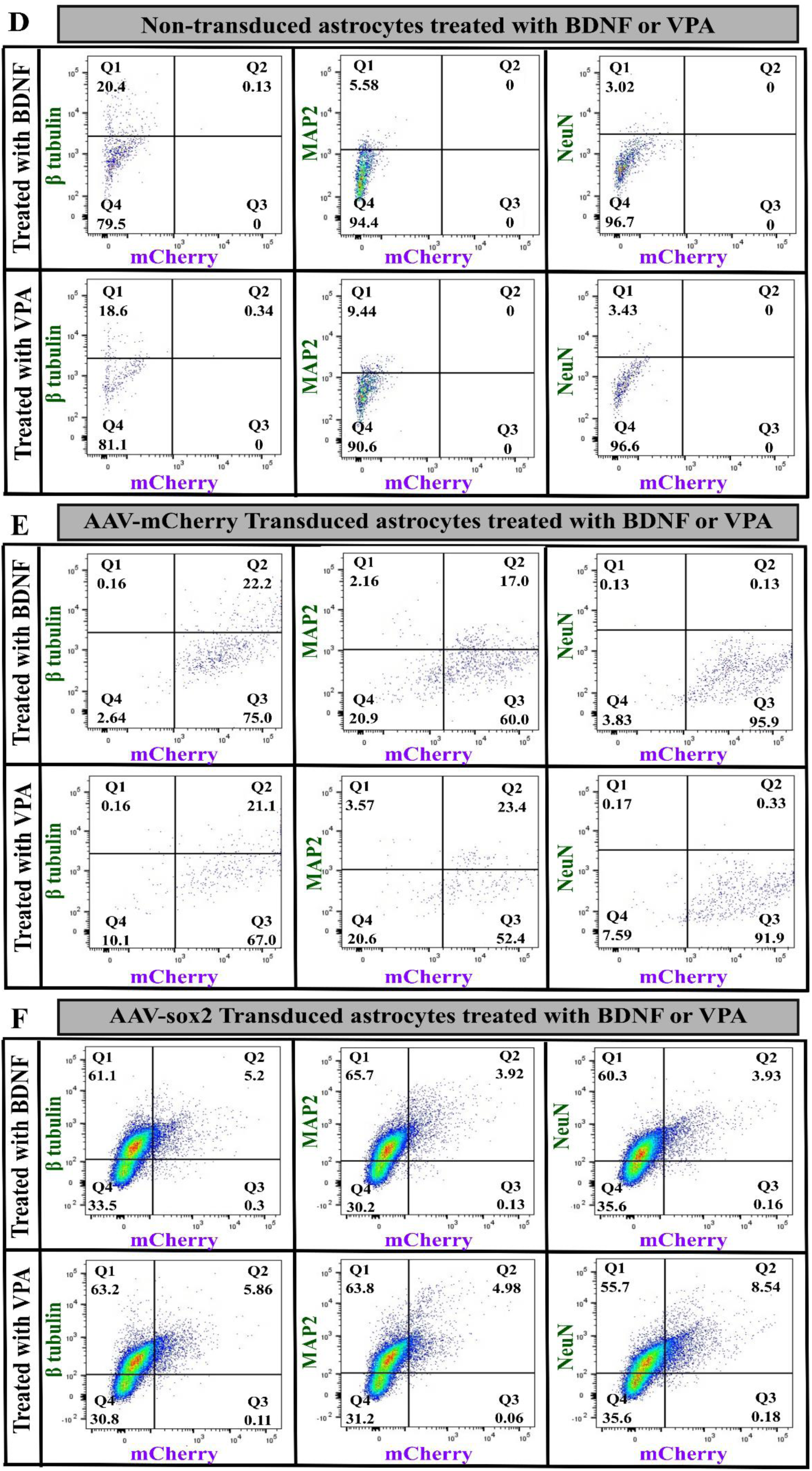
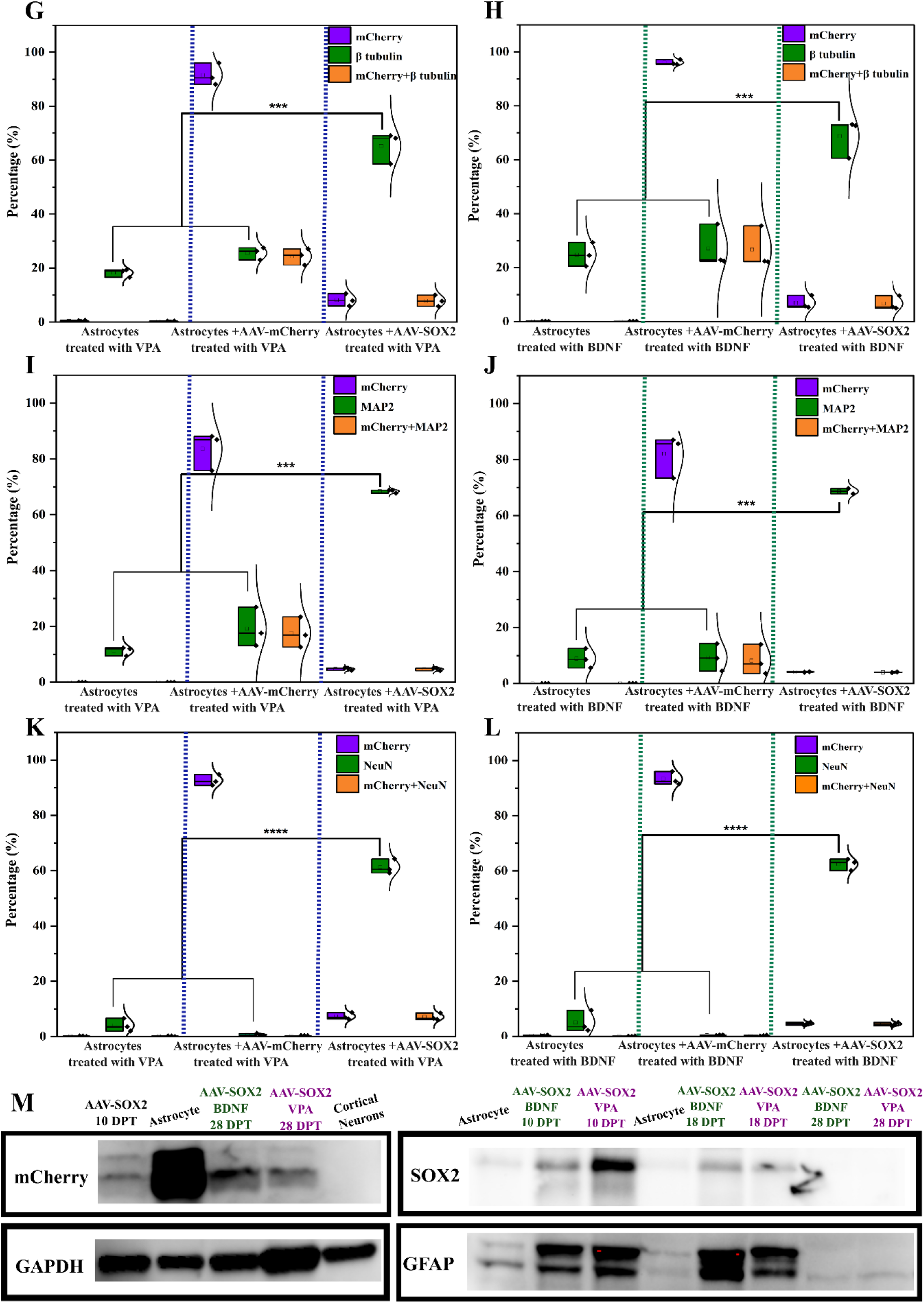
Morphological changes of primary mouse astrocytes in non-transduced/ AAV-mCherry transduced/ or AAV-SOX2-mCherry transduced astrocytes supplemented/treated with either BDNF/or VPA *in vitro*. A representative image of **A)** non-transduced primary astrocytes, **(B)** Empty vector (AAV-mCherry) transduced astrocytes, and **(C)** AAV-SOX2-mCherry transduced astrocytes treated with BDNF/or VPA for 28 days and stained with β3-tubulin/ MAP2/or NeuN (green), mCherry (red), GFAP (yellow), and DAPI (blue). **D-F**) Representative flow cytometry plots for non-transduced/ AAV-mCherry/ and AAV-SOX2-mCherry transduced astrocytes reveal different population of positive cells for different markers. **G-L)** Quantified cell counts for different cell markers with different conditions at 28 DPT. Scale bar= 100 µm. *** *p < 0.001*, **** *p < 0.0001*. n=3/group. **M)** Western blotting analysis of mCherry, SOX2, and GFAP protein level in AAV-SOX2 (treated with BDNF or VPA), AAV-mCherry transduced astrocytes at different time points of reprogramming as well as a positive control (cortical neurons). GAPDH served as control.

Using AAV-SOX2-mCherry, we were able to produce cells expressing β3-tubulin+, MAP2+, or NeuN+ neuronal markers with high reprogramming efficiency *in vitro* (>50%)^51^. These cells with neuronal fate are expected to be in post-mitotic stages. To investigate this, cell cycle analysis was performed on AAV-SOX2 and AAV-mCherry transduced astrocytes at 28 DPT. As seen in (**Figure S2A**), both S and G1 phases were significantly different between AAV-mCherry **(Figure S2D, Di)** and AAV-SOX2-mCherry transduced astrocytes after 28 DPT (**Figure S2Diii**).

Given that the conversion of astrocytes to neuron-like cells was confirmed through immunohistochemistry, using cortical primary neurons as a control would be more appropriate to determine whether the cells have reached a post-mitotic stage. As demonstrated in **Figure S2B**, the G2/M phase is significantly different between AAV-SOX2-mCherry transduced cells at 28 DPT compared to cortical neurons as control (**Figure S2Dii**). The difference between the G2/M phase in control and AAV-SOX2- mCherry transduced cells may be due to SOX2 transduction giving rise to astrocytes and oligodendrocytes as well as neurons. Thus, to selectively study neuronal-like cells, only the NeuN+ expressing cells were examined for their cell cycle stages. The results (**Figure S2C**) confirmed no statistically significant difference between the cell cycle of NeuN+ cells in AAV-SOX2-mCherry transduced cells (**Figure S2Div**) and cortical neurons, which suggests the reprogrammed iN cells are in a post-mitotic state and therefore not tumorigenic.

Moreover, the dynamic expression of SOX2 during the reprogramming process is important since at the early stage, high expression of SOX2 is required for initiating endogenous multipotency gene expression^53^. On the other hand, the persistent expression of SOX2 results in the maintenance of progenitor characteristics and hinders reprogramming to neurons^54^. Therefore, the subsequent downregulation of SOX2 is important for cells to exit cell cycle and undergo terminal differentiation. The ability to modify expression of SOX2 can be achieved through gene expression under the hGFAP promoter which is high in astrocytes but rarely expressed in neuronal cells^52,48,49^.

As illustrated in **Figure S2E,** reprogramming of astrocytes to neurons using AAV-SOX2-mCherry resulted in a drastic reduction in mCherry+ cells. This is likely attributed to the downregulation of SOX2 and gfaABC1D promoter during the reprogramming process to give rise to neurons. Moreover, during the M phase, many TFs and chromatin binding proteins are ejected from the chromatin; the integrative viral vectors such as lentiviral vectors are gradually silenced, and the non-integrative/episomal viral vectors such as AAVs are gradually removed from host cells. In addition to mCherry downregulation, at 28 DPT the population of cells expressing appropriate GFAP+ also drastically decreased in AAV-SOX2-mCherry transduced cells, further reinforcing the fact that the newly generated neurons are the result of the conversion of astrocytes and are not occasional preexisting neurons that may still reside in astrocyte culture. Moreover, 7.5 ± 2.1% of cells were double positive for β3-tubulin and GFAP marker demonstrating the portion of cells still in transitioning state after 28 DPT (**Figure S2F)**.

To further demonstrate the different protein expression level changes (GFAP, SOX2, mCherry) over time during reprogramming, AAV-SOX2-mCherry transduced astrocytes were collected at different time points to perform western blot. As expected, reprogramming of astrocytes to neurons resulted in down-regulation of astrocytes (GFAP signal), mCherry (AAV signal), and SOX2 (AAV and progeny marker signal) **(Figure 3M)** over time. After 28 DPT, the results demonstrate the absence of SOX2, mCherry, and GFAP signal which further confirms our observation through fluorescence microscopy.

Although both BDNF and VPA showed similar improvements in reprogramming efficiency, the *in vivo* use of BDNF is hindered by its susceptibility to enzyme degradation and short half-life^55^. Several studies have reported the capability of VPA to i) reduce HDAC activity and promote neuronal differentiation and outgrowth of NSCs^5657^, ii) upregulate the expression of diverse neurotrophic factors^58^, iii) facilitate white matter repair and neurogenesis after a stroke^59^, iv) exert an anti-inflammatory effect^60^, v) reduce cellular apoptosis and death of motor neurons and ^61^, vi) decrease demyelination and axonal loss, protects oligodendrocytes and neurons^62^, vii) reduce cystic cavitation^63^, and viii) ability to regulate chromatin accessibility and improve transduction efficiency^64,65^. More importantly, VPA facilitates shifting towards neuronal fate and inhibits glial fate simultaneously through the induction of neurogenic transcription factors including NeuroD1^56^. Consequently, VPA was selected to be embedded in electrospun nanofibers for subsequent *in vivo* application.

### Reprogrammed Primary Astrocytes are Able to Initiate Action Potentials

To further clarify the fate of the reprogrammed cells and confirm the generation of functional neurons from reactive astrocytes, we assessed the cells’ ability to generate action potentials using electrophysiological recordings^66,67^. Whole-cell patch-clamp results at 28 DPT showed that AAV-SOX2-mCherry transduced neuronal cells rarely become functionally mature and only exhibited spikelets (**Figure S3)**. Moreover, the resting membrane potential (RMP) of - 60 mV in both VPA and BDNF treated cells demonstrates partial neuronal maturation (**Figure S3A,I**). Thus, to achieve mature neurons with neuron identity, cells were assessed at more protracted periods after AAV-SOX2-mCherry transduction.

By 42 DPT, the recorded cells displayed typical mature RMP (*i.e.*, approximately between - 60 and -80 mV; **Figure 4A, J**) in which spontaneous calcium activity was also observed, indicating intrinsic excitability often observed in neurons (**Figure 4B, K**). At 42 DPT, current-clamp membrane depolarizing steps elicited action potentials in AAV-SOX2-mCherry transduced cells treated with VPA (n=5 cells) (**Figure 4C, D**) and BDNF (n=6 cells) **(Figure 4I, J).** Cells also displayed typical action potential (AP) kinetics, including a threshold near - 20 to -30 mV **(Figure 4E, N),** AP amplitude of 30 mV or larger **(Figure 4G, P)**, as well as typical rise and decay times (*i.e.*, <2 ms rise; **Figure 4H, I, Q, R**) observed in neuronal cells. The faster rise and decay time observed in reprogrammed cells at 42 DPT compared to 28 DPT indicates better sodium and potassium channel kinetics as expected in cells with neuronal identity. Slight differences observed in membrane potential and action potential kinetics such as action potential threshold might reflect the gradual maturation process of reprogrammed neurons. Together, these data suggested that by 42 DPT, SOX2 TF was capable of reprogramming astrocytes into functional neurons. We have culture primary cortical neurons as a positive control to see at which stage are the reprogrammed cells. Cortical neurons were patched at different time points 2, 5, 14, and 21 days in vitro (div). As shown in Figure **S4A-H**, at 2div no action potential was observed in response to current injections with the patched cells with RMP of -40 mV, confirming the presence of non-excitable/ immature neurons. At 5div, the cells were able to fire either single or repetitive action potential (**Figure S4I-P**) in response to current injections, with RMP of around -40 mV. As presented in **Figure 4**, in the reprogrammed cells at 42 DPT cells were able to fire action potentials in response to current injections but with more negative RMP, indicating the reprogrammed cells have developed sufficiently to establish a hyperpolarized resting state compared to cortical neurons at 5div. We have also recorded the activity of cortical neurons at 14div **(Figure S5A-H)** and 21div **(Figure S5I-P)** in which cells were able to fire repetitive action potential with RMP of -60 mV which reveals the presence of fully functional neurons at this stage. These results suggest that reprogrammed cells at 42 DPT show intrinsic electrophysiological properties similar to primary cortical neurons between 5div and 14div.

**Figure 4.**
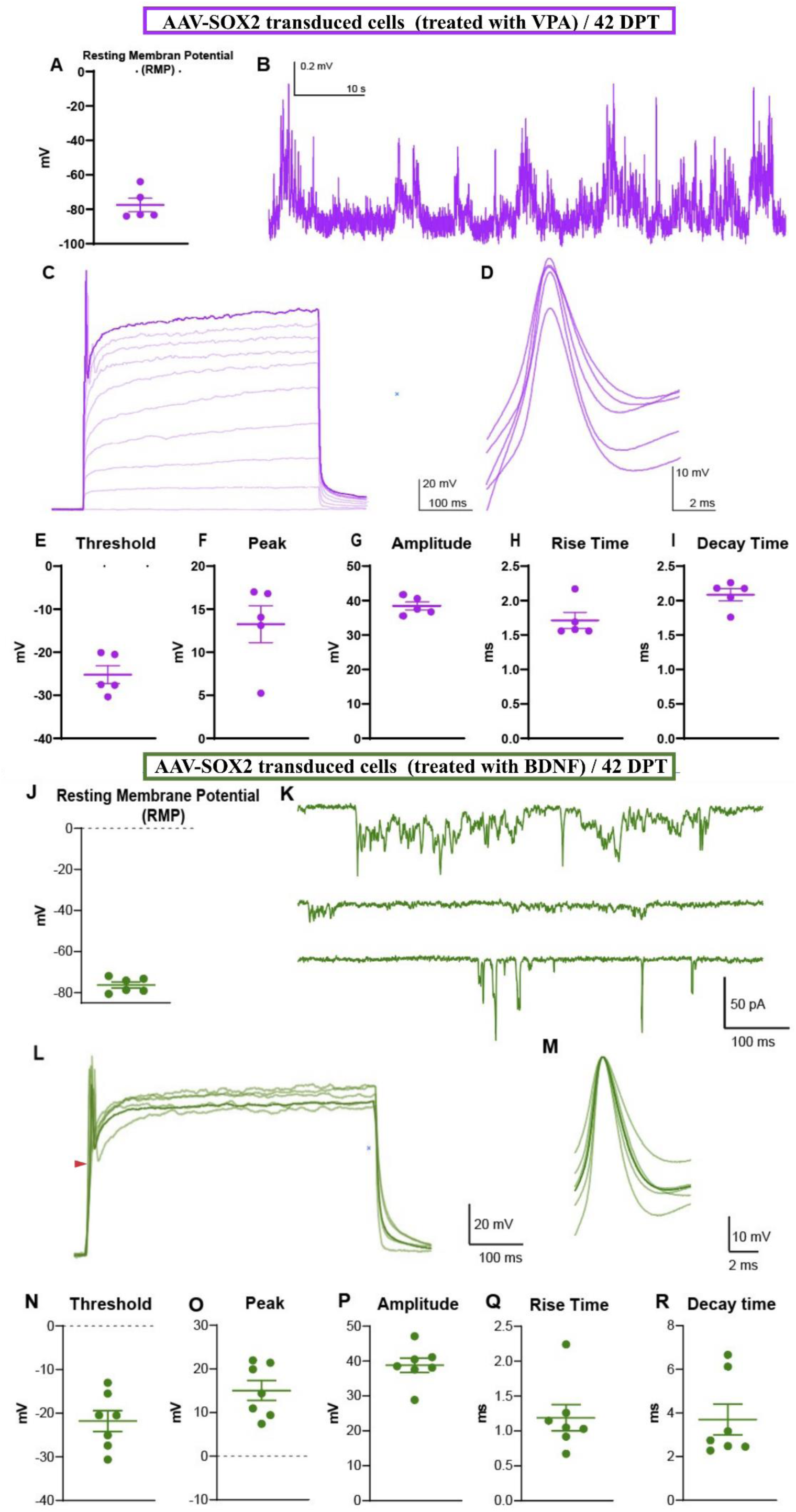
The SOX2-mediated reprogramming in both VPA and BDNF supplemented media was monitored *in vitro* and functionally characterized. **(A, J)** Resting membrane potential (RMP) for SOX2 transduced cells supplemented with VPA (n = 5 cells) and BDNF (n = 6 cells), respectively resembling RMP of mature neurons. **(B, K)** Representative examples of the spontaneous activity in recorded cells (B: current clamp at RMP; K: voltage clamp, -70 mV holding potential). **(C, L)** Cell depolarization and action potential resulting from current injection (C: overlaid incremental depolarization and appearance of an action potential in a single cell across 100 pA current steps; L: overlaid depolarization and action potential presentation across cells). **(D, M)** Overlaid action potentials (APs) from all the recorded cells. **(E, N)** AP threshold that regulates the flow of information which is less negative than RMP. **(F, O)** Peak potential of AP. **(G, P)** Amplitude of AP (from threshold to peak) reflecting the integrity of functioning axons. **(H, Q)** Rise time of AP (10%-90% of amplitude). **(I, R)** Decay time of AP.

### Assessment of Hybrid Composite Biomaterial

Satisfied we had a robust and controlled methodology for the reprogramming of astrocytes into mature neurons, we designed a hydrogel composite using the requirements of the molecular biology to ensure a bespoke, effective delivery system. Electrospun nanofibers offer high surface area and broad flexibility, while hydrogels provide highly controlled properties, including 3D networks, biodegradability, and mechanical strength^68^. Due to the unique properties of hydrogel and electrospun nanofiber, there has been increasing interest in combining these polymers for various biological and biomedical fields^69–72^. Electrospinning can be used with natural or synthetic scaffold materials and can accommodate a wide range of drugs, including therapeutic agents, proteins, growth factors. For example, it has been shown that this composite scaffold can provide a sustained, continuous release of growth factor for 28 to 90 days without affecting the bioactivity confirmed by neurite outgrowth from PC12 cells *in vitro*^73–75^. In our study, we loaded VPA into polylactic acid (PLA) electrospun nanofibers to provide a controlled and sustained release of the payload over time. Then, the fabricated electrospun scaffold was cut into short fibers and mixed into a Fmoc-DDIKVAV SAP hydrogel, called hybrid composite biomaterial, to develop a material with the capacity to be injected in the site of injury, fill the void, and provide a compatible interface with adjacent brain tissue.

### Hybrid Composite Biomaterial Prolongs the Delivery of VPA

PLA ± VPA electrospun nanofibers were formed using applied voltage. Scanning electron microscopy (SEM) imaging was employed to assess and confirm the morphology and alignments of the resultant fibers. Both groups of electrospun nanofibers (PLA ± VPA) exhibited a smooth, bead-free homogeneous structure, indicating the optimal flow rate and polymer concentration (10%)^76^, with uniform diameters **(Figure 5A, B)**. The inclusion of VPA in the electrospinning solution did not adversely affect the shape, morphology, and porosity of the resultant electrospun fibers **(Figure 5A, B)**. Image analysis revealed that the average fiber diameter of fabricated electrospun nanofibers was 1.6 ± 0.9 µm **(Figure 5C)**. Subsequently, PLA ± VPA electrospun nanofibers were cut to 20 μm lengths using a Leica microtome to make short fibers (SFs). The SEM images and image analysis of the resultant SFs (PLA ± VPA) displayed the homogenous fiber distribution **(Figure 5D, E)** with length of 22.4 ± 7.4 µm and 28.4 ± 8.2 µm for PLA SFs and PLA + VPA SFs, respectively **(Figure 5F)**.

**Figure 5.**
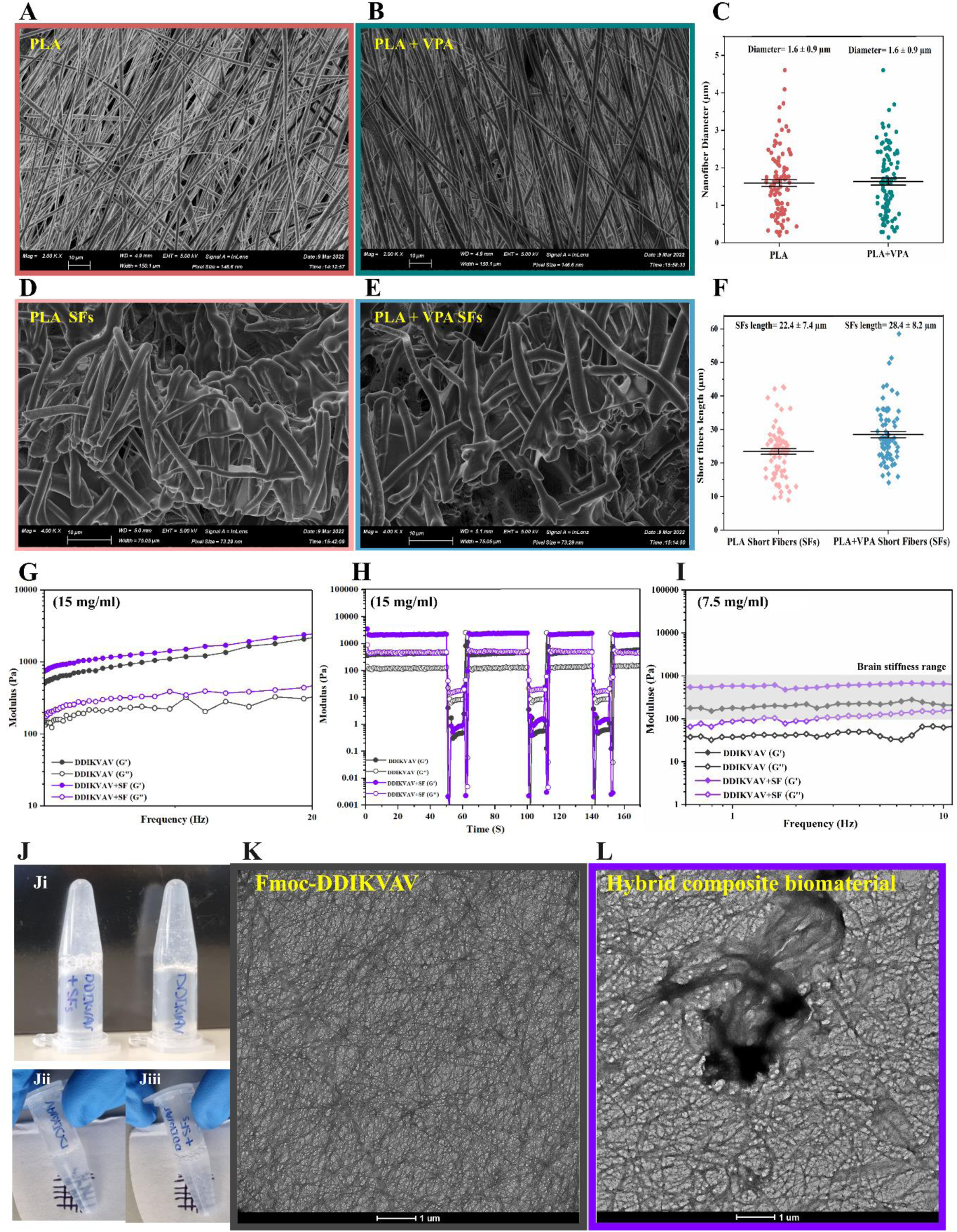
Fabricating the optimal hybrid composite biomaterial with encapsulated VPA. SEM images illustrating the nanofiber structure of the **A)** PLA electrospun nanofiber, **B)** PLA + VPA electrospun nanofiber, **C)** The corresponding distributions of the PLA ± VPA, **D)** PLA short fibers (SFs), **E)** PLA + VPA SFs, and **F)** The corresponding distributions of the PLA ± VPA SFs based on measuring about 100 fibers from the images. Scale bars= 10 µm. **G)** Assessment of the viscoelastic properties of the Fmoc-DDIKVAV SAP hydrogel and hybrid composite biomaterial showing higher G’ (storage modulus) than G” (loss modulus), **H)** Shear-thinning properties of SAP hydrogel and hybrid composite biomaterial, **I)** The viscoelastic properties of the Fmoc-DDIKVAV SAP hydrogel and hybrid composite biomaterial at 7.5 mg/ml concentration showing stiffness within brain’s stiffness range 100-100 Pa, **J)** Optical images of the fabricated hydrogels indicating Ji) inversion test and transparency of (Jii) SAP hydrogel and (Jiii) hybrid composite biomaterial. **K, L)** TEM images showing the nanofiber structure and network integrity of the SAP hydrogel (K) without and (L) with SFs incorporation to fabricate the hybrid composite biomaterial. Scale bars=: 1 µm.

The average diameter and length of the fibers was determined using ImageJ software by measuring about 100 fibers from the images obtained by SEM. No significant statistical difference was observed between the SF length in PLA ± VPA (one-way *ANOVA,* p<0.05).

Microenvironment stiffness is a critical factor influencing stem cell growth, differentiation and cell-reprogramming process^77,78^. Cells can sense the stiffness of ECM, convert these extracellular biophysical cues into intracellular biochemical signals through mechanisms known as mechanosensing and mechanotransduction. This process regulates chromatin organization, gene expression, protein translation, and overall cell function in time dependent manner^79–81^. To examine this, substrate stiffness/elasticity and shear-thinning analysis of the Fmoc-DDIKVAV SAP hydrogel and subsequent hybrid composite biomaterial were studied to make sure structure and viscoelastic behaviour of hydrogels maintained after the incorporation of SFs. Both fabricated Fmoc-DDIKVAV SAP hydrogel and hybrid composite biomaterial (15 mg/mL) and (7.5 mg/mL) demonstrated viscoelastic behaviour (G’>G”) **(Figure 5G,I)**. The composite biomaterial with the concentration of 7.5 mg/mL exhibited stiffness within the range of the brain (100-1000 Pa) and was selected for the subsequent *in vivo* study (**Figure 5I**). Moreover, the shear-thinning properties of the SAP hydrogels and the hybrid composite biomaterial were examined to confirm the self-recovery of the hydrogels after injection which is an important factor affecting payload outflow **(Figure 5H)**. The results demonstrate that the inclusion of SFs in SAP hydrogel did not disrupt the viscoelastic behaviour and slightly improved the stiffness of the hydrogels which is completely in line with other studies showing the inclusion of electrospun nanofibers reinforce the mechanical properties (higher modulus) compared to pure hydrogel^82,83^. Optical images of both fabricated SAP hydrogel and hybrid composite biomaterial confirmed the stable **(Figure 5Ji)** and clear **(Figure 5Jii, Jiii)** 3D gel formation. To further study the structure of the fabricated materials, transmission electron microscopy (TEM) imaging was performed **(Figure 5K, L)**. As shown in **(Figure 5L)** SFs were encapsulated within the structure of the SAP nanofibers without disrupting their integrity.

We examined the release kinetics of the VPA from hybrid composite biomaterial *in vitro* **(Figure S6A)**. Upon casting hybrid composite biomaterial, PBS was added on top and release profile of VPA was evaluated by analysis of supernatant at defined intervals followed by PBS replenishment. LC-MS analysis indicated that the hybrid composite biomaterial provided prolonged release of VPA over 8 weeks **(Figure S6B)**. These findings demonstrate that VPA encapsulated within the hybrid composite biomaterial will provide long-term delivery.

### Incorporating AAV-SOX2-mCherry within a SAP Hydrogel Sustains Viral Delivery while Retains Viral Functional Activity

To demonstrate the controlled release and functional activity of the AAV, the AAV-SOX2-mCherry was mixed with Fmoc-DDIKVAV SAP hydrogel before gelation (SAP + AAV) then cast in 96 well plates and equilibrated in an incubator overnight. Then to determine the release profile of AAV, PBS was collected at defined time points and analyzed using qPCR **(Figure S6C)**. Analysis revealed the sustained release of the AAV-SOX2-mCherry from SAP hydrogel **(Figure S6D)**. Moreover, to confirm the functional activity of AAV-SOX2-mCherry, after casting SAP + AAV in 24 well plates and equilibration overnight, astrocytes were cultured on top of the SAP + AAV, changing the medium every 2-3 days. Cells were monitored for 10 days after transduction. As seen in **(Figure S6E),** AAV-SOX2-mCherry transduction gave rise to colonies of small cells resembling NSCs confirming cells underwent proliferating states while in control AAV-mCherry transduced cells this behavior was not observed **(Figure S6F)**.

### SOX2 Forced Expression Generates Intermediate Neuroblast Cells

To determine our ability to drive neurogenesis in the rodent brain, AAV-SOX2-mCherry was delivered ectopically to the host striatum. Neurogenesis around the injection site was examined by staining for the expression of doublecortin (DCX). It has been reported that neurogenesis through SOX2 ectopic expression is a slow process and DCX+ cells are not readily detectable until 28 DPT^54^. Our observation on 28 DPT confirmed the generation of DCX+ cells mainly surrounding the AAV-SOX2-mCherry injection site **(Figure S7A)**. In sharp contrast, DCX+ cells are sparsely detected along or surrounding the site injected with empty vector (AAV-mCherry), indicating the absence of neuroblast cells **(Figure S7B)**. It has also been reported that, continual expression of SOX2 using CAG promoter induced approximately 37-fold fewer DCX+ cells compared to dynamic expression of SOX2 provided by hGFAP promoter^52^. These data indicate that the constitutive expression of SOX2-mCherry is not necessary and might even be detrimental to induced adult neuroblast cells (intermediated cells) expansion and survival. Here we observed that the induced DCX+ cells also did not express mCherry marker confirming the dynamic expression of SOX2 and silencing during the process of reprogramming (**Figure S7C**). Moreover, it has been reported that the newly generated DCX+ neuroblast cells can eventually generate mature neurons when supplied with VPA^84^. Therefore, to examine that we have developed a hybrid composite biomaterial to provide sustained release of VPA to the site of injury.

The phenotype and function of astrocytes and microglia are strongly influenced by the CNS microenvironment. Under normal conditions or even in the unaffected areas of a diseased CNS, microglia exhibit a normal phenotype with appropriate density^85,86^. It has been shown that AAV can disrupt the BBB and induce immune cell infiltration in a titer-dependent manner, even in intracranial injection which is considered as a relatively safe procedure^87^. Herein, we have studied different field of views (FOVs) perpendicular to needle track (**Figure 6A**) and observed elevated density of reactive astrocytes and microglia in the AAV-mCherry + PBS-VPA as well as AAV-SOX2-mCherry + PBS-VPA group (high-titer AAV area), whereas the density of these inflammatory cells was significantly attenuated in the AAV-SOX2-mCherry + SAP-SF-VPA group, suggestive of the presence of the conducive microenvironment provided by the hybrid composite biomaterial and enhanced repair process (Figure S8). Our approach resulted in less reactivity of astrocytes and microglia which may, in the long term, facilitate brain recovery.

**Figure 6.**
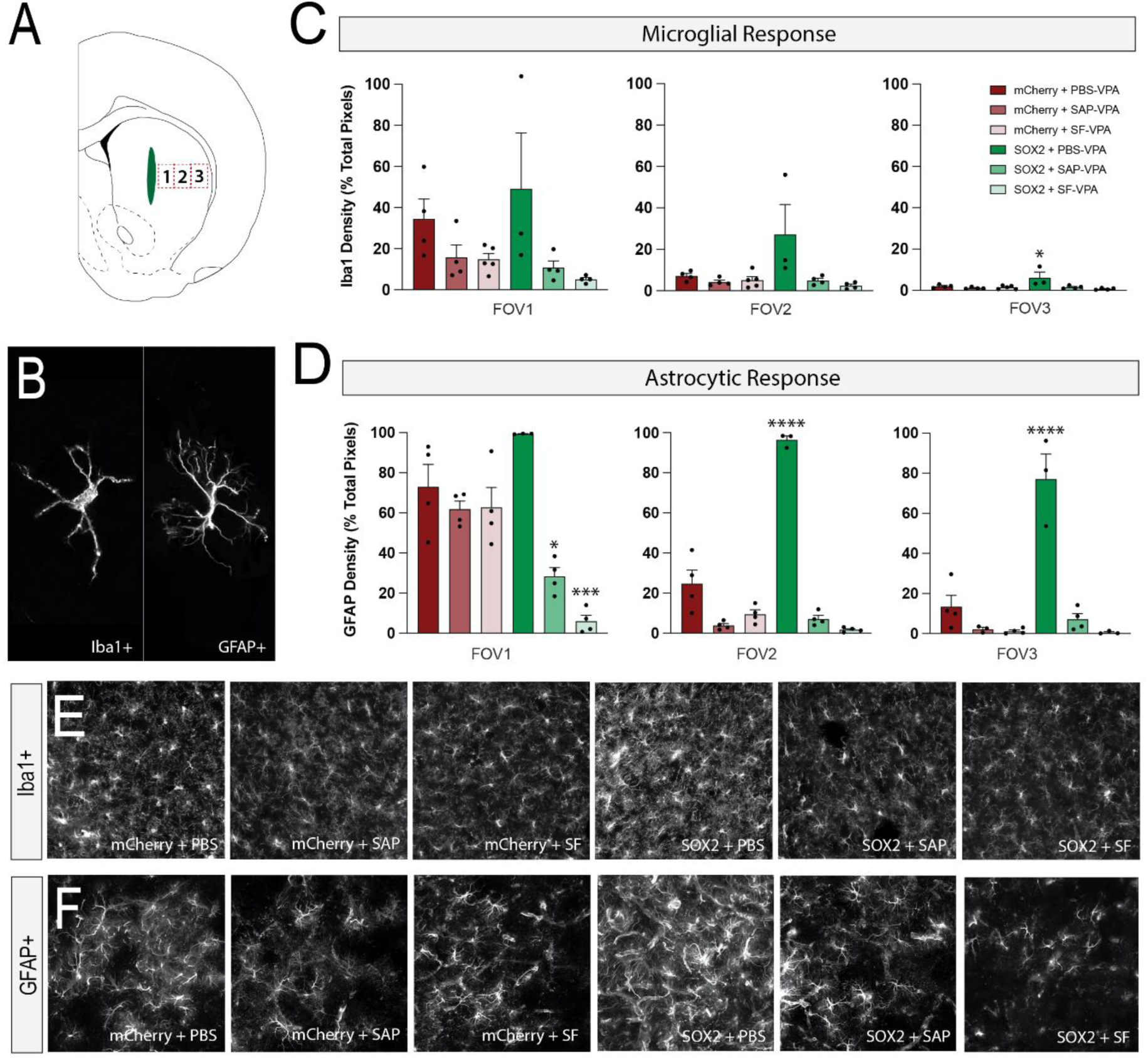
The synergistic effect of AAV-SOX2-mCherry and composite biomaterial reduced glial scars and microglial inflammatory responses. **A)** Schematic diagram of the different field of view (FOV) selected for analysing the density of reactive astrocytes and microglial cells, **B)** Higher magnification of single cell of reactive microglia (Iba-1) and astrocyte (GFAP). Quantified measurements of the density of **(C)** microglia and **(D)** astrocytes. Representative fluorescent images of the coronal sections (FOV3) of different groups stained for **E)** Iba-1 and **F)** astrocytes. *p < 0.05, ****p < 0.0001 vs relative control. Data are represented as the mean ± SEM (n = 4−5 per group). Scale bar= 100 µm

Apart from the morphological changes to microglia and astrocytes after injury, the activated glia polarizes in response to different factors towards different phenotypes^88^. The inflammatory cascade activated after brain trauma is mediated by the release of pro- and anti-inflammatory cytokines which are rapidly upregulated in response to pathological conditions^89^. Moreover, cytokines function as mediators of intracellular communication and play a pivotal role in tissue homeostasis^89,90^.

The inflammatory cytokines produced by leukocytes, astrocytes, microglia, and neurons induce local and systemic immune responses^91^. It has been shown that the pro-inflammatory phenotype releases deleterious factors (e.g., tumour necrotic factor alpha (TNF-𝛼), interlukine-6 (IL-6), IL-1β, interferon gamma (IFN-𝛾), MMPs, reactive oxygen species (ROS), and etc) leading to tissue damage and inflammation^92^. These pro-inflammatory cytokines promote the reactive phenotype of macrophages (M1). The microglia activation can induce A1-responsive astrocytes^92^. On the other hand, an anti-inflammatory phenotype that secretes neurotrophic factors (e.g., IL-3, IL-10, transforming growth factor beta (TGF-β), VEGF, BDNF, NT-3, IGF-1, GLT-1) acts as a neuroprotectant and induces reparative astrocytes (A2) to help suppressing inflammatory response^92^.

Herein, we have analysed different cytokines to further examine the changes in anti- and pro-inflammatory factors in different groups. To do so, the PFA-fixed tissues were de-crosslinked and extracted proteins examined using cytokine spotted array membranes (**Figure 7 A and B-G**). Hierarchical clustering (Euclidean, average linkage) of the cytokines confirmed the distinct secretome signatures between different treatments. The dendrograms (**Figure 7H**) indicate that the AAV-SOX2 + SF and AAV-SOX2 + SAP have similar secretome profile. As shown in **Figure 7H**, delivery of AAV-SOX2-mCherry with the hybrid biomaterials resulted in an increased level of anti-inflammatory cytokines such as IL-3 and IL-10 and a reduction in the pro-inflammatory cytokines such as IL-1α, IL-1β, TNF-α and IFN-𝛾.

**Figure 7.**
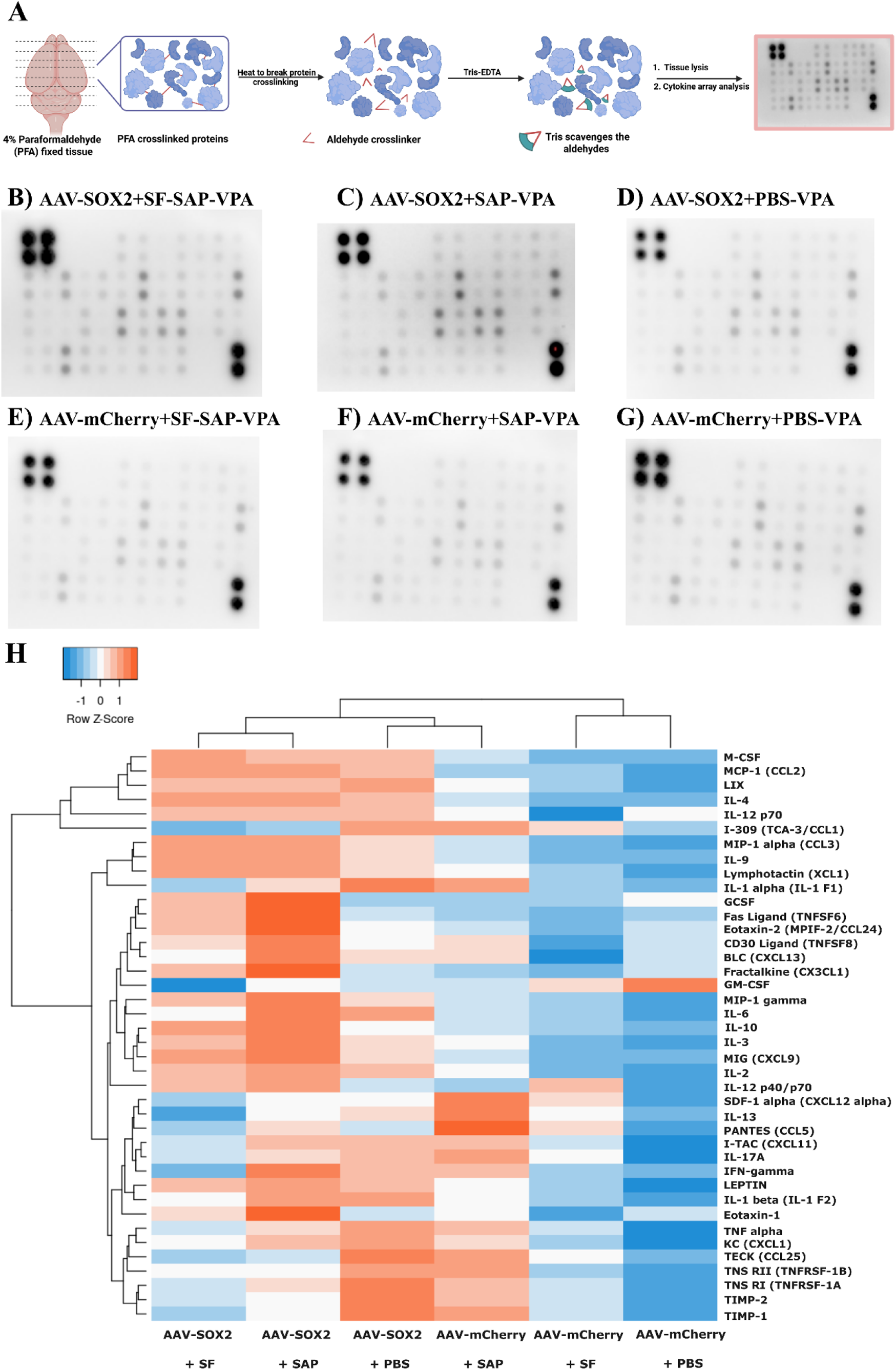
Heatmap and dendrogram showing clustering of samples with similar cytokine expression pattern patterns in six experimental groups. **A)** Schematic of the procedure for protein extraction from paraformaldehyde (PFA) fixed tissue, created with BioRender. The photographs of cytokine arrays in which each cytokine is represented by duplicate spots arrays for 40 different cytokine probes on a single membrane for **B)** AAV-SOX2-mCherry + SAP-SF-VPA, **C) B)** AAV-SOX2-mCherry + SAP-VPA, **D)** AAV-SOX2-mCherry + PBS-VPA, **E)** AAV-mCherry + SAP-SF-VPA, **F)** AAV-mCherry + SAP-VPA, and **G)** AAV-mCherry + PBS-VPA. **H)** Heatmap hierarchical clustering representation of the mean expression of the 40 inflammatory proteins (semi-quantitative) in different groups. Orange color indicates an increased cytokine production compared to the control, whereas a blue color indicates the suppression in the production of the cytokine.

IFN-𝛾 can induce the activation of microglia to M1 phenotype which results in the secretion of numerous pro-inflammatory mediators^93^. Moreover, IL-6 which is highly elevated in AAV-SOX2-mCherry is due to the higher level of pro-inflammatory microglia. In contrast, fractalkine (CXC3CL1), IL-4, and IL-13 can trigger a transformation in microglia toward M2 phenotype^94^. Based on **Figure 7H** the elevation of IL-4 and fractalkine is observed in AAV-SOX2-mCherry + materials. The cytokine expression results are in line with the morphological changes of astrocytes and microglia as discussed in **Figure 6**. Together, the cytokine expression levels and morphological changes in presence of material confirms the importance of ECM-mimicking material to modulate inflammatory response and potentially aid any recovery.

### Intermediate Neuroblast Cells Mature into Neurons

The cellular fate of proliferating intermediate neuroblast cells (iNBCs) was evaluated by labelling with bromodeoxyuridine (BrdU) and the neuron-specific nuclear protein NeuN. We tested the effect of sustained release of VPA on the maturation of the iNBCs by treating mice twice daily with 50 mg kg^−1^ of BrdU i.p. injection for 2 weeks (beginning 2 weeks post AAV/material implantation). Then we examined the results on the stained tissues of mice 8 weeks post implantation. As shown in **Figure 8A, B** differentiation of intermediate cells into mature neurons were confirmed with cells double stained for BrdU+ and NeuN+, indicated with arrowhead. The mature neurons, derived from intermediate cells, were observed more in the striatum of mice injected with AAV-SOX2-mCherry + hybrid composite biomaterial (SOX2 + SAP-SF-VPA) **(Figure 8B)**. This observation could be attributed to the sustained release of VPA that diffuses out of the SFs into a hydrogel and then into the surrounding tissue. In addition, the initial release of VPA from either SAP hydrogel or hybrid composite biomaterial could be functional for the treatment of TBI, contributing to the reduction of the cascade of secondary events and attenuation of a specific cellular response. As seen in **Figure 6** and **Figure S8**, the AAV delivery in different groups resulted in varying levels of astrocyte activation. In groups with elevated astrocytes reactivity the number of cells as a source for reprogramming is higher than the groups with less astrocytes. Consequently, the number of new neurons (NeuN+BrdU+) was normalized to the proportion of GFAP+ reactive astrocytes **(Figure 8D)**.

**Figure 8.**
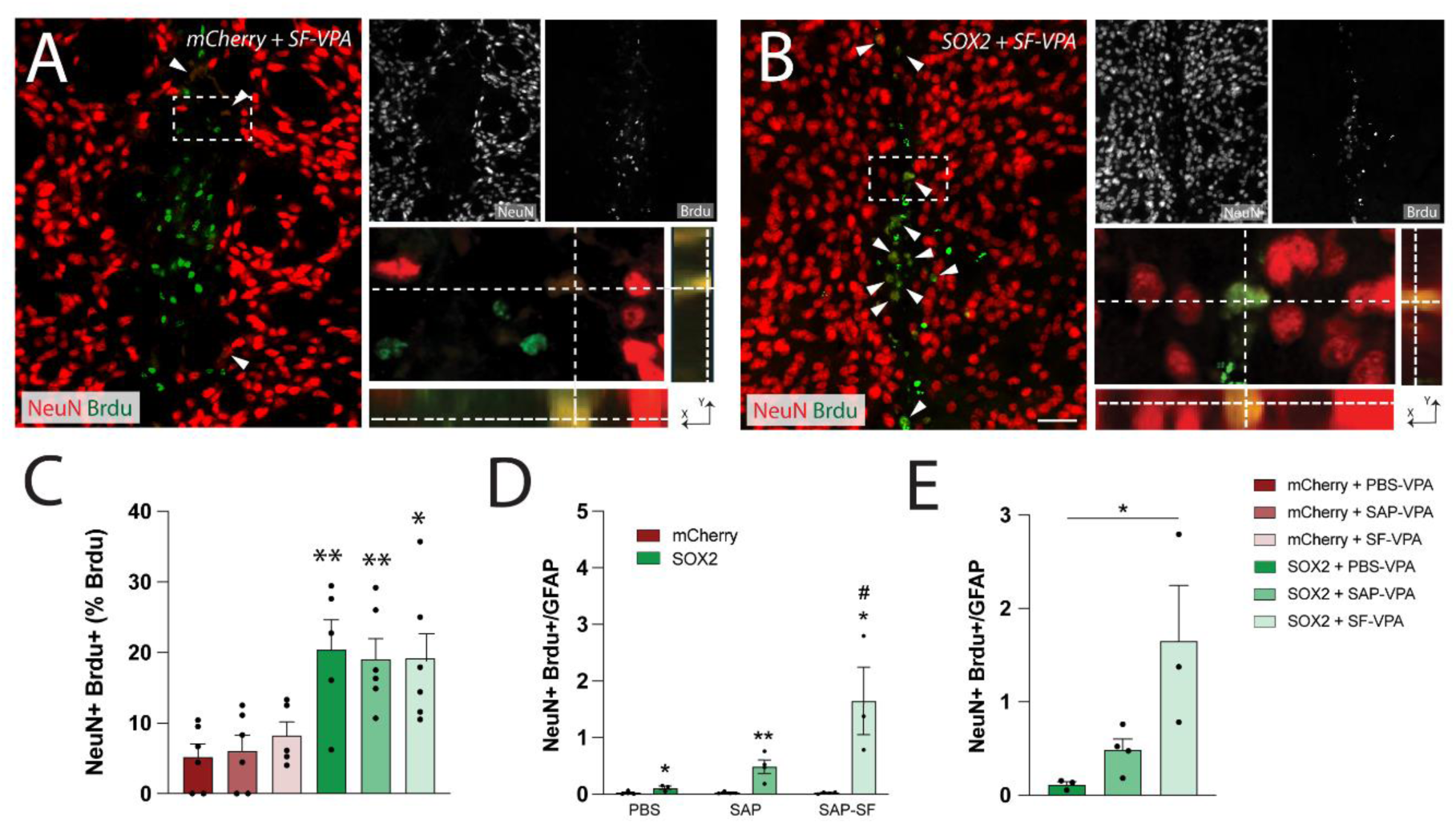
Representative photomicrographs of A) AAV-mCherry + SAP-SF-VPA and B) AAV-SOX2-mCherry + SAP-SF-VPA labelled positive for both BrdU and NeuN markers. Arrows show some representative cells stained positive for both NeuN and BrdU. **C)** treatment with AAV-SOX2-mCherry significantly increased the proportion of BrdU+ NeuN+ in the site of injection. **D)** NeuN+Brdu+/GFAP density demonstrates a significant increase in SOX2 groups compared to their relevant mCherry controls. **E)** Interestingly, when compared AAV-SOX2-mCherry + SF showed significantly higher NeuN+Brdu+/GFAP compared with AAV-SOX2-mCherry + SAP and AAV-SOX2-mCherry + PBS, demonstrating the higher reprogramming efficiency in the presence of hybrid composite biomaterial and sustained release of VPA. *p < 0.05, **p < 0.01 vs relative controls (D). Data are represented as the mean ± SEM. n=4-5/group. Scale bar= 100 µm

Here we confirmed that the ectopic expression of SOX2 robustly induced iNBCs, followed by the generation of mature neurons in the presence of VPA diffused from hybrid composite biomaterial. This approach resulted in a significant increase of BrdU+ cells being labelled by NeuN+ in AAV-SOX2 + SF-VPA treated mice during 2 weeks’ time **(Figure 8)**.

Our ability to generate mature neurons from the pool of iNBCs resulting from the dedifferentiation of resident astrocytes with a single dose injection of materials together with AAV-SOX2 is in the pursuit of a potential regeneration-based therapies for CNS-associated injuries or diseases.

## Conclusion

Despite extensive research efforts, regenerating damaged areas of the central nervous system (CNS) after injury remains a significant clinical challenge. Reprogramming large numbers of proliferating glia, including astrocytes, represents an innovative approach to replenish neurons lost in response to neural insult or degeneration.

In this study, we present a composite biomaterial that closely mimics the brain microenvironment and serves as a controlled-release platform for reprogramming factors. We validated the effectiveness and efficiency of the SOX2 transcription factor in reprogramming reactive astrocytes into induced progenitor neural cells (iPNSCs) and subsequently into mature neurons. This reprogramming was facilitated by the sustained release of valproic acid (VPA) from the composite material. The synergistic effect of the SOX2 TF and VPA incorporated within the hybrid composite biomaterial significantly enhanced the efficiency and safety of the epigenetic conversion of astrocytes to mature neurons. Our findings present a promising avenue for translating this approach into regenerative medicine applications for CNS repair.

## Materials and methods

### Polylactic acid (PLA) Scaffold Fabrication, Valproic Acid (VPA) Immobilization, and Characterization

#### PLA Nanofiber Scaffold Fabrication

Electrospinning is a technique to produce electrospun fibers by employing electrostatic forces. This appealing method produces fine fibers with diameters ranging from several micrometers to a few nanometers depending on the polymer types and processing conditions^95^. Herein, to fabricate electrospun nanofibers, PLA (Polylactic acid) was obtained from GoodFellow. Polymer solutions of 10% (w/v) were made with chloroform and acetone (3:1 v/v) and magnetically stirred at room temperature (RT) to dissolve completely. For valproic acid (VPA) loaded fibers, VPA was mixed with the PLA solution at the ratio of 1/30 (v/v).

The prepared solution was then loaded into a plastic syringe with an 18G×11/2 gauge needle, and programmable syringe pump. The solution was dispensed at the rate of 2 mL h^-1^ while charged to 20 kV at a working distance of 15 cm. The fibers were collected on an aluminium collector, rotating at 1000 rpm during a 2.5-hour period.

### Short Fiber (SF) Production

Electrospun nanofibers (PLA ± VPA) were cut at the size of 20 μm on a Leica microtome (SM 2010R). Dry ice was crushed and used to freeze small pieces of the scaffolds (2 × 1 cm) in an upright position embedded in OCT cryo-mounting media (TissueTek OCT, Fisher Scientific, USA). After cutting, the resultant SFs were collected and washed 3 times with deionized water (DI) to remove the OCT and then recovered by centrifugation at 4000 rpm for 15 min.

### Solid Phase Peptide Synthesis (SPPS), SAP Hydrogel/and Hybrid Composite bioMaterial Preparation

The Fmoc-DDIKVAV peptide was fabricated using solid-phase peptide synthesis (SPPS) method, as previously described^96^. The resultant peptide powder was used to fabricate the self-assembling peptide (SAP) hydrogels through pH-switching method. Briefly, 10 mg of the peptide powder was suspended in 400 µL DI H_2_O and was fully dissolved by the minimal addition of 0.5 M sodium hydroxide (NaOH). Then, self-assembly was induced by dropwise addition of 0.1 M hydrochloric acid (HCl), while vortexing. PBS was then added to achieve the final peptide concentration of 15 mg mL^-1^. To fabricate hybrid composite biomaterial (Fmoc-DDIKVAV + SFs), SFs were mixed with SAP hydrogel prior to gelation. For *in vivo* and *in vitro* cell culture, Fmoc-DDIKVAV (95-99% desalted) was manufactured by Pepmic, China and the fabricated gel was exposed to ultraviolet (UV) light at least for 20 min.

### Rheology

The rheological analysis was performed using an Anton Paar Rheometer in a cone plate geometry. 200 µL of the sample, SAP hydrogel alone and hybrid composite biomaterial, was placed in the middle of the plate and allowed for 5 min to set before testing to stabilize. Rheological analysis was performed with a gap size of 0.102 mm, multiple frequency sweeps with frequencies ranging from 0.1-100 Hz, and a 0.1% oscillatory strain at a constant required temperature (37 °C).

### Scanning Electron Microscopy (SEM)

Scanning electron microscopy (SEM) was performed on a Zeiss UltraPlus electron microscope employing a 5.0 kV beam and an in-lens detector. Briefly, electrospun nanofibers ± VPA and SF ± VPA as solution droplets (10 µL) was added on a double-sided carbon tape sputter coated with Platinum for 60 seconds at 20 mA. After imaging with SEM, nanofiber diameter and length were analysed using ImageJ.

### Transmission Electron Microscopy (TEM)

Negative stain TEM analysis was conducted using a Hitachi H7100FA with a tungsten filament fitting at 100 kV. Approximately 4 μL of hydrogel/ hybrid composite biomaterial was added on top of the glow-discharged (15 mA, 30 s) Formvar-coated copper grids for ≈ 1 min and immersed in DI H2O twice for washing. The grids were then flipped on 10 µL of uranyl acetate (UA, 1%) as contrast-enhancing agent solution for ≈ 1 min. Excess liquid was blotted off after every step using Whatman paper. Then the grids were left to air dry overnight before TEM imaging.

### Composite Hybrid Biomaterial Fabrication and VPA Release Profile Investigation

To investigate VPA release profile, 150 µL of the hybrid composite biomaterial casted in 96 well plates (n=3) and placed in incubator (37 °C, 5% CO_2_) to equilibrate. After 1h, 100 µL of PBS was gently added on top. At appropriate intervals of 1, 8, 24 h, and 1, 2, 3, 4, 5, 6, 7, 8 weeks the supernatant was collected and replenished with an identical volume of fresh PBS. The collected samples were stored at −20 °C until analyses with LC-MS.

### Release profile *in vitro* Using High Performance Liquid Chromatography (HPLC)-Orbitrap

The VPA concentrations were determined by LC-MS spectrometer (Thermo Scientific Q-Exactive Focus hybrid quadrupole-Orbitrap). 40 µL of collected supernatant from composite biomaterial, were transferred to HPLC vials and dried down using SpeedVac (ambient temperature/1 h), then dispersed in 100 µL of Milli Q water + 0.1% Formic acid (FA), vortexed and sonicated, filtered through a 0.2-µm membrane filter, and centrifuged at 1000 rpm for 5 min. A total of 10 μL of each sample was injected (flow rate of 0.2 mL min^−1^) into LC-MS spectrometer (Thermo Scientific Q-Exactive Focus hybrid quadrupole-Orbitrap). Chromatographic separation was achieved using an ACQuity C18, LC column of 2.1×50 mm, 1.7 µm. The mobile phase was a mixture of 0.1% FA in water and acetonitrile 95:5 (v/v).

### AAV Vector Production and Purification

AAV production was performed at the Vector and Genome Engineering Facility (VEGF) at the Children’s Medical Research Institute (CMRI), Sydney, Australia. Briefly, HEK293 cells (ATCC) were used as a substrate for AAV production using PEI (polyethylenimine MW 25000), with a 1:1:2 molar ratio of pCap:pTransgene:pAd5. Vector particles purification was performed using standard iodixanol gradients (4 different layers of gradient (15%, 25%, 40%, and 60%) containing PBS-MgKNa, Optiprep, and Phenol Red (just for 25% and 60% gradient)). Vector particles were collected from a thin layer between 40%-60% gradient buffer. Amicon Ultra-4 Centrifuge Filter Units with Ultracel-100 kDa membrane (EMD Millipore, Cat# Z648043) were used for final buffer exchange (Phosphate-buffered saline [PBS, Gibco, Cat# 14190], 50 mM NaCl [Gibco, Cat#24740], 0.001 %, Pluronic F68 [v/v] [Gibco, Cat# 24040]). Vector particles were collected, precipitated, and concentrated by centrifugation and concentration step (volume 200-500 μl). Vector preparations were quantified using droplet digital PCR (ddPCR) (qPCR, Bio-Rad, Cat# 172-5125) using mCherry primers (F: 5’-CACTACGACGCTGAGGTCAA-3’; R: 5’-GTGGGAGGTGATGTCCAACT -3’). In this technique the sample is partitioned into thousands of tiny individual droplets in which the PCR amplification occurs. The fraction of positive droplets was then used to calculate the absolute concentration of the constructed AAV. We have achieved the of 1.02×10^13^ vg mL^−1^ and 9.6×10^12^ vg mL^−1^ for AAV-DJ gfaABC1D-SOX2-mCherry (AAV-SOX2) and AAV-DJ-gfaABC1D-mCherry (AAV-mCherry), respectively.

### Primary Culture

#### Primary Astrocytes

Primary astrocytes were isolated from the cortices of the brain of postnatal day (P)1.5-2 Swiss mice pups as previously described^18,46^. Briefly, the cortices of the brain were dissected in the cold DMEM High Glucose media. After meninges removal the forebrain tissues were mechanically dissociated to generate a single-cell suspension. Subsequently, cells were centrifuged at 1000 rpm for 10 min and resuspended in the complete DMEM (CDMEM) 10:10:1 culture medium containing (DMEM, 10% fetal bovine serum (FBS) (Gibco), 10% horse serum (HS) (Gibco), and 1% penicillin/streptomycin (Hyclone)), transferred to a poly-D-lysine (PDL) coated 75 cm^2^ flask, and incubated at 37 °C.

When cultures became confluent (14 days’ post culture), non-astrocytic cells were removed from the cell monolayer by shaking vigorously overnight on an orbital shaker (230 rpm at 37 °C). Then the medium was collected as a cell source for astrocytic and non-astrocytic cells to examine whether the fabricated AAVs are more prone to transduce astrocytic cells or non-astrocytic cells.

To collect astrocytes, the flask containing attached astrocytic cells Then, the medium was discarded, and the flask was gently washed 3 times with PBS (1X) for the complete removal of the medium and non-astrocytic cells. Cells were then dissociation with 3 mL trypsin/EDTA (10 min at 37 °C) pelleted for 10 min at 1000 rpm and resuspended in the fresh astrocytic medium composed of CDMEM 5:5:1.

### Primary Cortical Neurons

To isolate primary cortical neurons, Swiss mice were time mated overnight (as the embryonic day E 0.5). Then, E14.5 fetuses were collected from the pregnant mouse. After removing the olfactory bulb and meninges, the collected cortices were dissociated into a single-cell suspension through trypsinization (digestion in HBSS with 1X 0.25% trypsin (Gibco), 1X DNAse (Gibco) for 20 min at 37 °C) and mechanical dissociation in serum-free minimum essential medium (MEM) solution with 10% FBS. Then cells were loaded with desired density into PDL coated culture plates and allowed to attach for 3 hours. Then the medium was replaced with fresh primary neurobasal culture media (neurobasal medium (Gibco) with 2% serum-free B27 supplement (Gibco), 5 mM glutamine (Gibco) and 10 μg mL^-1^ gentamicin (Sigma)).

### Astrocytic Cell Transduction/Transfection

The primary astrocyte cells were used to evaluate the transduction efficiency of AAV-SOX2. For this purpose, astrocytes were loaded into 24-well plates with PDL coated glass coverslip (Thermo Fisher, Waltham, MA, USA) at a density of 2 × 10^4^ cells per well. To transduce astrocytes, the AAVs were thawed on ice, followed by the addition of an appropriate amount of AAV-SOX2 into each well, plus negative control wells (multiplicity of transduction (MOT=0)), and positive control AAV-mCherry (MOT=5) and incubated for 24 h. Then to minimize viral exposure, the media was completely replaced with a fresh differentiation medium. After 10/ 28/ 42-day post transduction (DPT), the transduced cells were fixed and immunostained to examine the efficiency of transduction and cell conversion.

### Protocol for Reprogramming, Neural Progenitor, and Differentiation culture

1 day after initial transduction, the medium was changed to the progenitor culture media: DMEM/F12 (Gibco) supplemented with 1× N2 and 1× B27, 20 ng mL^-1^ fibroblast growth factor (bFGF), 20 ng mL^-1^ epidermal growth factor (EGF), penicillin/streptomycin. For neuronal differentiation, media was supplemented with brain-derived neurotrophic factor (BDNF, 20 ng mL^-1^, Invitrogen)/ 0.5 µM valproic acid (VPA) to the cultured medium every 3-4 days to promote synaptic maturation during the differentiation.

### Conversion Efficiency, Neuronal Purity, and cell cycle analysis *in vitro*

The conversion efficiency and cell cycle were analysed by Flow cytometry (Fusion, BD Bioscience). Briefly, the adherent immunostained cells were gently detached from well plates using mini scraper and suspended in PBS. Then the conversion efficiency was calculated as the percentage of the number of β3-tubulin+, MAP2+, and NeuN+ cells to quantify reprogramming efficiency, mCherry+ cells to acquire transduction efficiency, and GFAP+ cells to study the population of astrocytes before and after reprogramming.

### Western Blotting

After transducing primary astrocytes with AAV-SOX2 and AAV-mCherry, cells were collected at different time points. The collected cell proteins were prepared with 1×cell lysis buffer containing RIPA buffer and Protease inhibitor. Then, the resulting protein concentrations were quantified with the BCA Protein Assays (Thermo Scientific, ZB382866).

The denatured proteins were loaded and separated with 12% Criterion TGX Stain-free Precast gel (BioRad) and transferred onto the PVDF membrane (BioRad, 1620177). The membrane was then blocked with 5% (w/v) fat-free milk in TBST (TBS with 0.1% Tween-20) for 60 min at room temperature, followed by incubation overnight at 4°C with the following primary antibodies: GFAP goat antibody (1:500, ab53554), mCherry chicken antibody (1:1000, ab205402), SOX2 mouse antibody (1:1000, ab171380), and GAPDH chicken antibody (1:1000, AB2302-25UG). The PVDF membrane was further washed with TBST and incubated with Donkey anti-Goat IgG (H+L) (1:1,000, 106-54-092524), Goat Anti-Mouse IgG HRP Conjugate (H+L) (1:1,000, 7105-40UL), and Donkey anti-Chicken IgY (H+L) (1:1,000, SA1-300) secondary antibodies for 1 h at room temperature. After washing with TBST, the Clarity Western ECL was added to each membrane and incubated for 2 min at RT, then the images were captured with Bio-Rad ChemiDoc Imaging System.

### Release Profiles

To determine the AAVs release profiles, 8 µL of AAV-SOX2 was incorporated within the Fmoc-DDIKVAV solutions prior to gelation (SAP-AAV solution). The SAP-AAV solutions were then self-assembled and transferred into a 96-well plate. After stabilization of hydrogel in incubator 100 µL of PBS was gently added to the top of the hydrogels and plates were incubated at 37 °C with 5% CO_2_. To determine AAV titers at different time points, PBS replenishment was conducted, and supernatants were collected and quantified using a qPCR AAV Titration (Titer) Kit (Abcam, Australia) as described before^18^. Briefly, the viral DNA, the collected supernatants were mixed with Virus Lysis Buffer (1:1 ratio) and incubated for 10 min at 70 °C to isolate the viral DNA. Then, 2.5 µL of the extracted viral DNA and AAV Standard 1 or AAV Standard 2 was mixed with 2X qPCR MasterMix (12.5 µL), Reagent Mix (10 µL) and Nuclease-free H_2_O (2.5 µL). Finally, the qPCR reaction was performed through the enzyme activation (10 mins at 95 °C), denaturation (95 °C, 40 times/15 secs duration each), and annealing/extension (60 °C, 40 times/1 mins duration each). Three technical replicates per time point were used.

### Electrophysiological Examination of SOX2 Converted Cells *in vitro*

After 28/42 DPT, the cell activity analysis of converted cells was examined using whole-cell patch clamp recordings. Cells on coverslip were transferred to the recording chamber then all recordings were obtained with a Multiclamp 700B (Axon Instruments) EPC9 amplifier and InstruTECH LIH 8+8 amplifier (Heka) at 21–23 °C. The standard ACSF external solution contained (in mM): 124 NaCl, 3 KCl, 2 CaCl_2_, 1 MgCl_2_, 1.25 NaH_2_PO_4_, 26 NaHCO_3_ and 10 glucose saturated with 95% O_2_ and 5% CO_2_, pH 7.2-7.4, and 275-285 mOsm. Whole-cell clamp recordings were performed using glass pipettes (5-8 MΩ) filled with the internal solution containing (in mM) 140 K gluconate, 10 HEPES, 2 NaCl, 4 KCl, 4 ATP, and 0.4 GTP at a pH of 7.4. Action potentials were elicited and recorded under the current-clamp mode via increasing steps of depolarizing current. Electrophysiological data was analysed in Easy Electrophysiology (RRID:SCR_021190). Action potential kinetics were determined using the automated detection modules, with the threshold calculated as the leading inflection (the minimum of the first derivative prior to the action potential peak).

### Electrophysiological Examination of Primary Cortical Neurons *in vitro*

After 2div and 5div, the cell activity analysis of primary cortical cells was examined using whole-cell patch clamp recordings. Cells on coverslip were transferred to the recording chamber then all recordings were obtained with a SEC-05X single electrode clamp amplifier and headstage (npi) at 21–23 °C. The recordings were digitised with 16-bit precision at 20 kHz (USB-6221, National Instruments). Instrument control and recording were performed using custom MATLAB scripts. The standard ACSF external solution contained (in mM): 124 NaCl, 3 KCl, 2 CaCl_2_, 1 MgCl_2_, 1.25 NaH_2_PO_4_, 26 NaHCO_3_ and 10 glucose saturated with 95% O_2_ and 5% CO_2_, pH 7.2-7.4. Whole-cell clamp recordings were performed using glass pipettes (5-12 MΩ) filled with the internal solution containing (in mM) 125 K-Gluconate, 2 CaCl_2_, 2 MgCl, 10 HEPES, 10 EGTA, 2 Mg-ATP and 0.5 Na_2_-GTP at a pH of 7.4. Action potentials were elicited and recorded under the current-clamp mode via increasing steps of depolarizing current (100 pA steps for 2div and 20 pA steps for 5div, 10 steps). Electrophysiological data was analysed using custom MATLAB scripts. The action potential threshold was calculated as the leading inflection (the minimum of the first derivative prior to the action potential peak). The rising time and decay time was determined as the 10%-90% time from threshold to peak.

#### In vivo

Adult female Swiss mice were used for this experiment. All procedures were performed in accordance with the Australian National Health and Medical Research Council’s published Code of Practice for the Use of Animals in Research and approved by The Florey Institute of Neuroscience and Mental Health Animal Ethics committee.

### Stereotaxic surgery and viral vector delivery

#### Needle stab injury

Mice were anaesthetised via 2% isoflurane inhalation and placed in a stereotaxic frame and using sterile procedures an incision made to expose the skull. A small burr hold was drilled into the skull through which a needle stick injury was made perpendicular to the exposed cortical surface using a Hamilton needle (external diameter 0.5 mm). Injury was made into the right and left striatum at the following coordinates: 1.0 mm anterior and +/-2.0 mm lateral from Bregma, and at a depth of 2.8 mm below the surface of the dura.

After 10 days, mice received bilateral microinjections of either (i) AAV-mCherry diluted in PBS-VPA, (ii) AAV-mCherry diluted in SAP-VPA; (iii) (i) AAV-mCherry diluted in SAP-SF; or (iv) AAV-SOX2 diluted in PBS-VPA; (v) AAV-SOX2 in SAP-VPA; (iv) AAV-SOX2 diluted in SAP-SF.

Note the dilution of AAV with PBS (or SAP/ SAP-SF) was 1:1 with 1µL/injection/hemisphere into the same site as the initial injury (AP-1.0 mm, L-2.0 mm, deep 2.8 mm below the surface of the dura), n=5-7 injections per group. The concentration of VPA in each group was 1 mM.

#### bromodeoxyuridine (BrdU) Administration

Dividing cells were fate-mapped by *in vivo* intraperitoneal injection (i.p.) of BrdU (B5002, Sigma; 50 mg kg−1 body weight in PBS, twice daily) for 2weeks starting at 14-days post AAV injection/material implantation. BrdU can be incorporated into the genome during the S phase of mitosis and detected when immunoreacted with an anti-BrdU antibody (555627, mouse, 1:500). Mice were killed 8 weeks post AAV/material implantation.

### Immunocytochemistry and Immunohistochemical Staining, Imaging and Analysis

For *in vitro* immunocytochemistry (ICC), cells were washed 3 times with PBS (1X) and fixed in 4% paraformaldehyde (PFA) solution for 20 min at RT, followed by 3 times PBS washes. Plates were then sealed with parafilm and stored in the fridge until immunocytochemistry.

To perform immunohistochemistry (IHC) on brain tissue sections, mice were killed by an overdose of sodium pentobarbitone (100 mg/kg) and transcardially perfused with Tyrode’s buffer, post-fixed in 4% PFA for 2h, and cryoprotected for 48 h in 30% sucrose. Afterwards, frozen brains were cryo-sectioned into 40-μm thick coronal sections using a microtome and serially collected (1:12) and stored in an anti-freezing solution at -20 °C.

Both IHC and ICC analysis were conducted as previously described. The following primary antibodies (neural and astrocyte markers) were used: mouse monoclonal anti-β3-tubulin/TUJ1, (1:500, biolegend), goat monoclonal anti-Glial Fibrillary Acidic Protein (GFAP, 1:500, Abcam), chicken polyclonal anti-mCherry (mCherry, 1:1000, Abcam), rabbit monoclonal anti-NeuN (Neun, 1:1000, Abcam)/ rabbit monoclonal anti-Microtubule-associated protein 2 (MAP2, 1:1000, Abcam), rabbit polyclonal anti-ki67 (1:300, Novus), and rabbit polyclonal anti-OCT3/4 (1:100, Invitrogen). Alexa Fluor-488/555/647 -conjugated corresponding secondary antibodies from Abcam (1:500) were used for indirect fluorescence. Samples were mounted, coverslipped, and visualized with either Zeiss 780 Confocal or Leica SP5 confocal microscope/Axioobserver at 10x, 20x, or 40x magnification with identical exposure settings. Additionally, to precisely analyse the rate of transduction and conversion efficiency for ICC, flow cytometry analysis was performed.

### Quantification

Astrocytic and microglial density in Figure 6 was measured as optical density at 3 defined fields of view (FOV) lateral of the needle stab injury (20x) of 1/12 series of chromogenic immunohistochemistry for GFAP and IBA1.

For *in vivo* BrdU+/NeuN+ cell counting in Figure 7, all BrdU+ cells were counted at the injury site in a 1/12 series, and the fraction of NeuN+ expressing cells colocalized with BrdU+ was quantified. Proportions were subsequently normalized to the density of GFAP reactivity **(Figure 8D/E).**

### Protein Extraction from PFA-fixed Brain Tissue

The PFA-fixed cryo-sectioned tissues in anti-freezing media were collected and washed 3 times with cold PBS. Then, to reverse the PFA crosslinks while maintaining the proteins, the tissue sections were heated to 80°C in Tris-EDTA solution (Antigen retrieval step).

After de-crosslinking PFA, the tissues were mechanically homogenized with a pestle in lysis buffer for 60 min on ice to aid solubilization. The tissue lysate was centrifuged at 14,000 x g, 15 min, 4°C. The supernatant containing the extracted protein was collected in a clean tube and kept at -80°C for further analysis.

### Cytokine Proteome Profiler Array

The inflammatory cytokines in the resected mice brains were analyzed using the Mouse Inflammation Antibody Array-Membrane (ab133999) according to the manufacturer’s instructions. Briefly, the array membranes were blocked with 1X Blocking Buffer provided in the kit. Then the tissue lysate with 200 µg of total protein was prepared in 1X Blocking Buffer and added to the array membranes following overnight incubation at 4°C. Then, the samples were aspirated from wells, followed by the large volume wash in Wash Buffer I, and then 3 times wash with 2 mL Wash Buffer I and 2 times with Wash Buffer II (5 mins) to minimize the background signals. The array membranes were then incubated with 1 mL 1X Biotin-Conjugated Anti-Cytokines for 1.5 - 2 h at RT on a platform shaker and washed with Wash Buffer I and II as described earlier. The membranes were further incubated with streptavidin-HPR in 1X Blocking Buffer for 2 h at room temperature before mixing with the Chemi reagent mix. Images were then captured with the Bio-Rad ChemiDoc Imaging System. ImageJ was used to quantify and determine spot density. To be able to compare results across different arrays, the signals of each biotin-conjugated spots were normalized to Positive Control signals. The Positive Control is like housekeeping in Western Blots.

## Statistical Analysis

Statistical tests employed (one-way ANOVA with Tukey’s post-hoc multiple comparison test and Student’s t-tests) are stated in the figure legends. *p < 0.05; ** p < 0.01; *** p < 0.001; **** p < 0.0001 indicated statistically significant difference. Data were reported as mean ± SD/SEM.

## Acknowledgement

DRN was supported by an ARC Future Fellowship. Funding: This work was funded by an National Health and Medical Research Support (NHMRC) Ideas Grant GNT 1144996 to D.R.N. and an Australian Research Council Discovery Grant DP220102549 to D.R.N. Access to the facilities of the Centre for Advanced Microscopy (CAM) with funding through the Australian Microscopy and Microanalysis Research Facility (AMMRF) is gratefully acknowledged. The authors thank Harpreet Vohra and Brianna Xuereb for their expert assistance.

## Data Availability Statement

The data that support the findings of this study are available from the corresponding authors upon reasonable request.

