## Supplementary material for "Spatially precise neuron formation via hydrogel mediated modulation of the host astrocyte response": SI

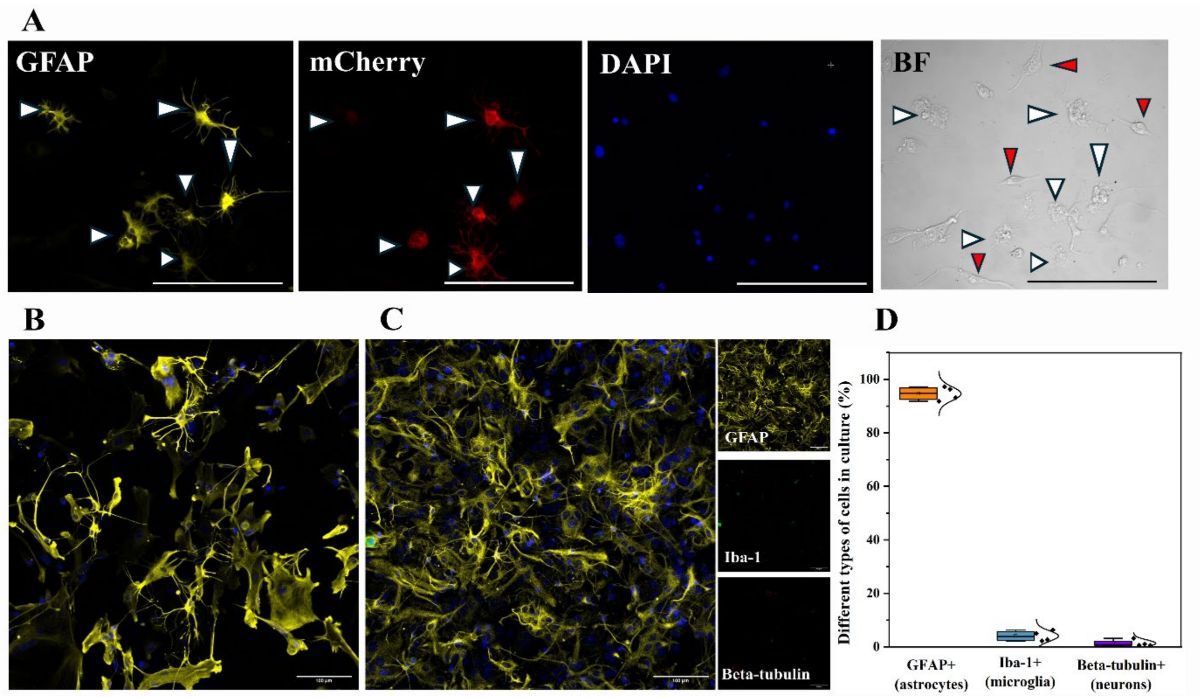


**Figure S*1*. *Specificity of the AAV construct to preferentially transduce astrocytes and purity of astrocytes in primary mouse astrocyte culture. A)*** *The mixture of different cell types (neuron, microglia, and astrocytes) transduced with AAV-mCherry and stained at 3 days post transduction (DPT) against transduction efficiency marker (mCherry, red), astrocytic protein GFAP (yellow), DAPI to study the transduction affinity toward different cell types. Some double stained cells with mCherry and GFAP are indicated with white arrowhead. Moreover, the red arrowhead in bright field image demonstrates the non-astrocytic cells in the culture which is not transduced with AAV-mCherry.* ***B, C)*** *To examine the purity of astrocytes, the primary astrocytes were stained against GFAP (yellow), Iba-1 (microglia marker, green), β-tubulin (neuronal marker, red), and DAPI.* ***D)*** Quantification data for GFAP+, Iba-1+, and β3-tubulin+ cells demonstrated the purity of astrocytes in culture (94.6 ± 2.5%). Data represents Mean ± SEM. *Scale bar= 100µm.*

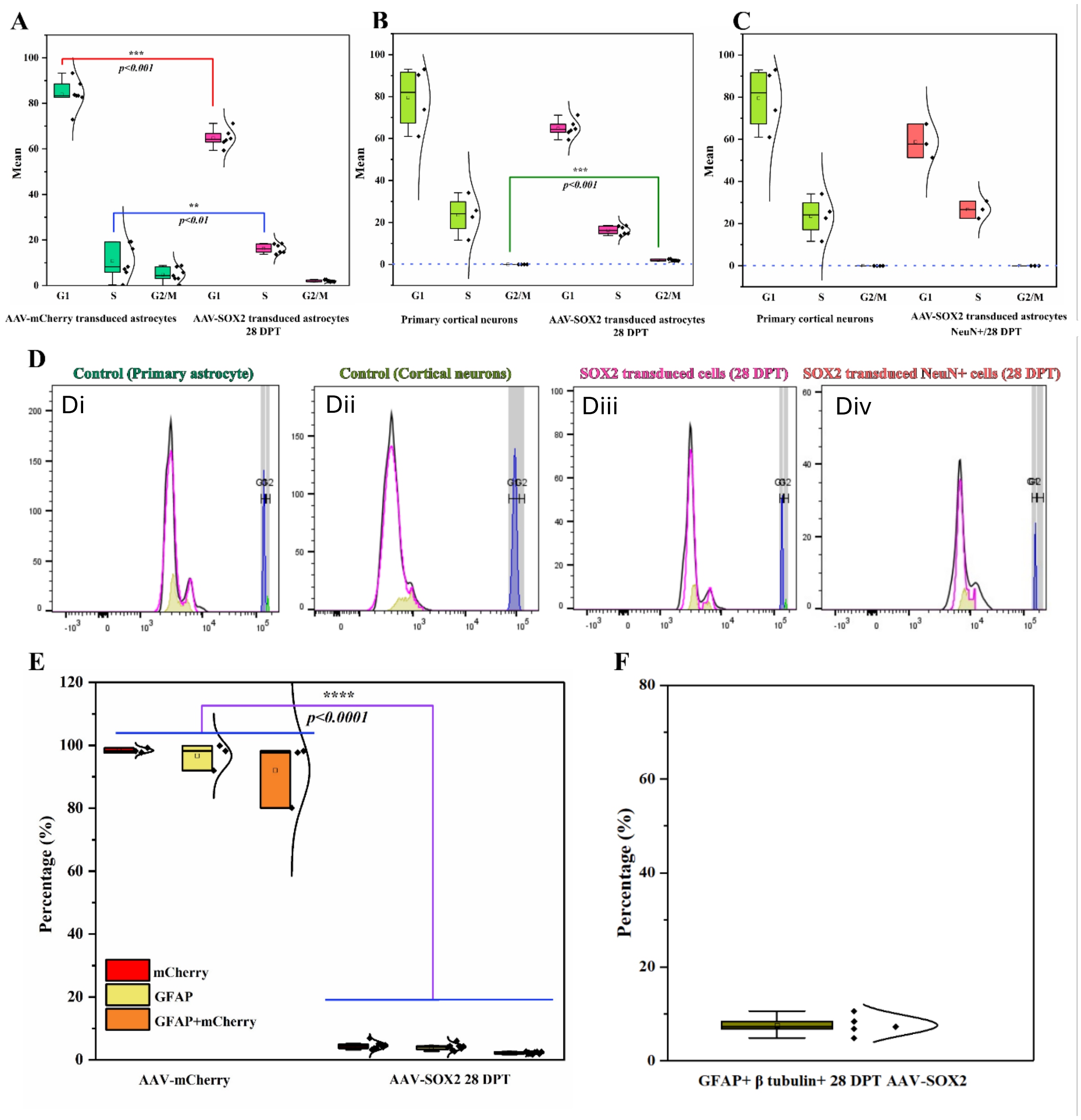


**Figure S2. Cell cycle analysis on transduced cells at 28 DPT**. **A-C)** Quantification of G1, S, and G2/M phases for primary astrocytes, primary cortical neurons, and AAV-SOX2 transduced astrocytes. **Di-Div)** Representative images of cell cycle plots acquired from flow cytometry for different cells. **E)** Quantification of GFAP+, mCherry+, and double positive cells (both GFAP+ and mCherry+), based on FACS data reveals a drastic reduction of mCherry+, GFAP+ and double positive cells after 28 DPT in AAV-SOX2 transduced astrocytes compared to AAV-mCherry transduced astrocytes. **F)** Quantification of β tubulin and GFAP positive cells 28 DPT, indicating the portion of cells in transitioning state. ** $p < 0.01$ *** $p < 0.001$, **** $p < 0.0001$. n=3/group

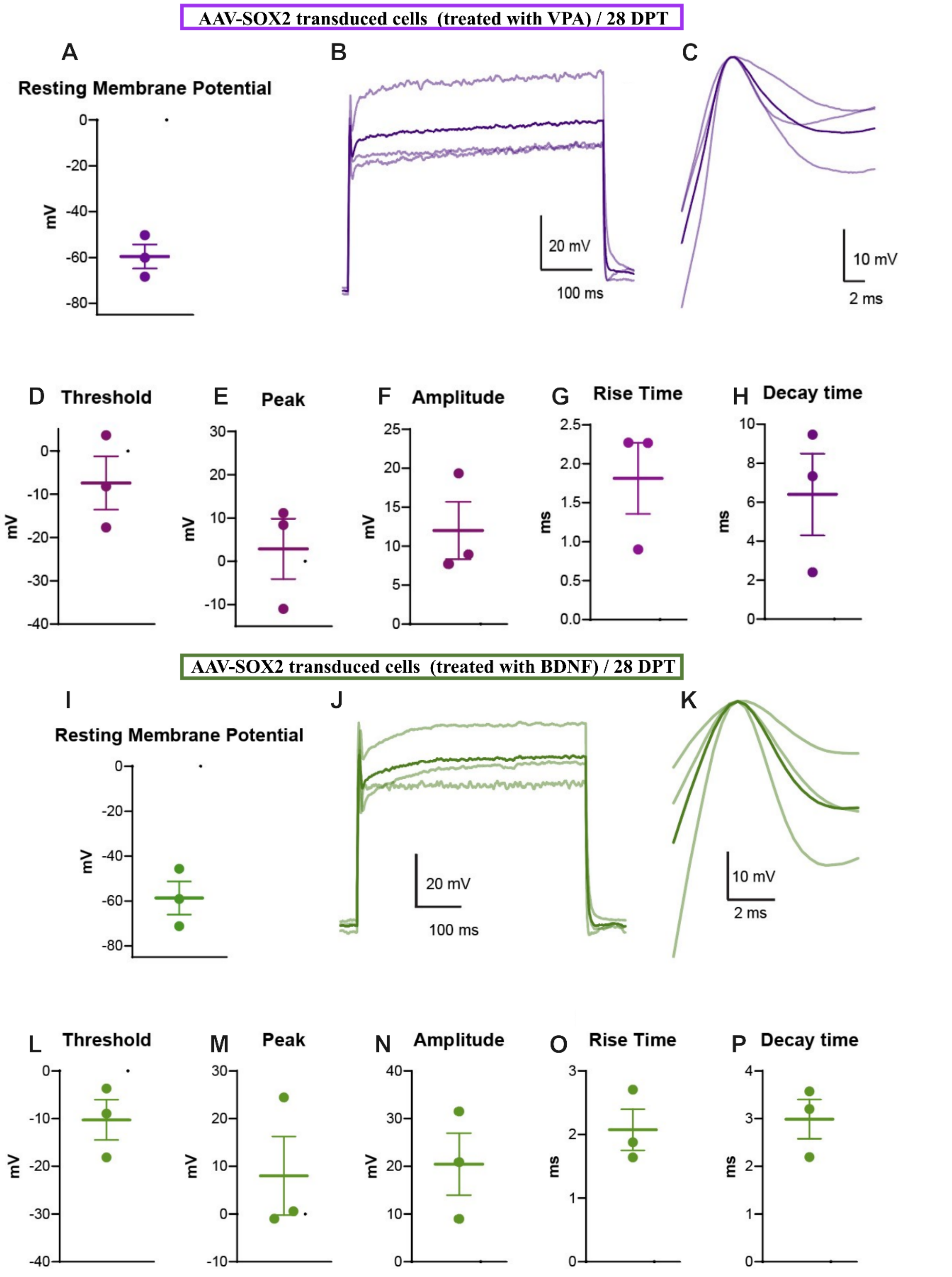


**Figure S3.** The SOX2-mediated reprogramming in both VPA and BDNF supplemented media was monitored *in vitro* and functionally characterized. **(A, I)** Resting membrane potential for SOX2 transduced cells supplemented with VPA (3 cells) and BDNF (3 cells), respectively. **(B, J)** Depolarization resulting from 25 pA current steps in cells. **(C, K)** Overlaid action potentials from all recorded cells. **(D, L)** Action potential threshold. **(E, M)** Peak potential of action potential. **(F, N)** Amplitude of action potential (from threshold to peak). **(G, O)** Rise time of action potential (10%-90% of amplitude). **(H, P)** Decay time of action potential.

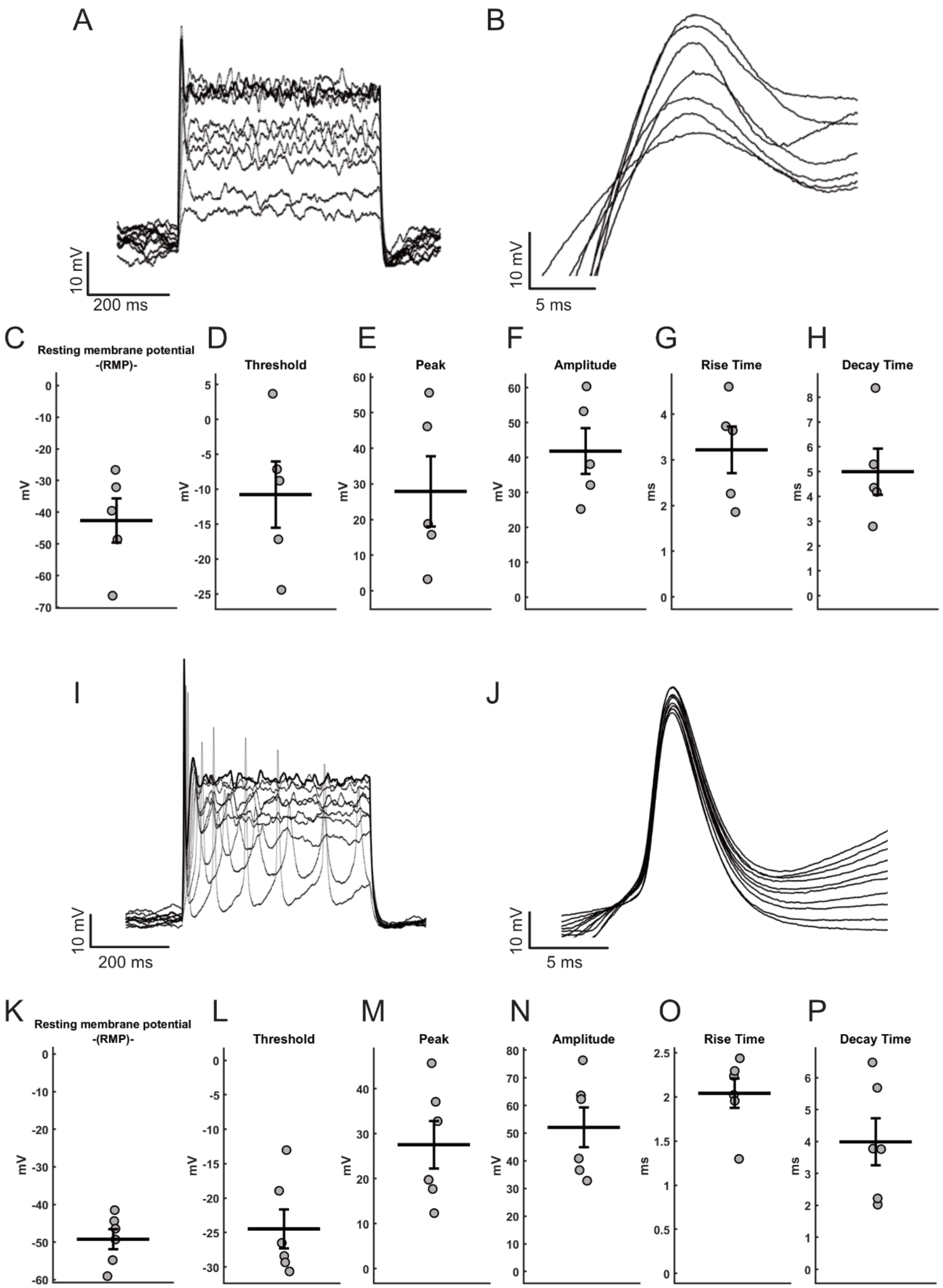


**Figure S4. Functional characterization of primary cortical neurons at timepoints of 2div (A-H) and 5div (I-P). (A, I)** Cell depolarization and action potential resulting from current injection (A: overlaid incremental depolarization and appearance of an action potential in a single cell across 100 pA current steps; I: overlaid incremental depolarization and appearance of an action potential in a single cell across 20 pA current steps). **(B, J)** Overlaid action potentials (APs) from all the events recorded during a single measurement. **(C, K)** Resting membrane potential for 2div (n = 5 cell) and 5div (n = 6 cells), respectively. **(D, L)** AP threshold. **(E, M)** Peak potential of AP. **(F, N)** Amplitude of AP (from threshold to peak). **(G, O)** Rise time of AP (10%-90% of amplitude). **(H, P)** Decay time of AP (90%-10% of amplitude). In graphs C-H and K-P the thick black line represents the average value over the population (2div, n= 5; 5div, n=6) while the grey dots represent the average value of the individual cells.

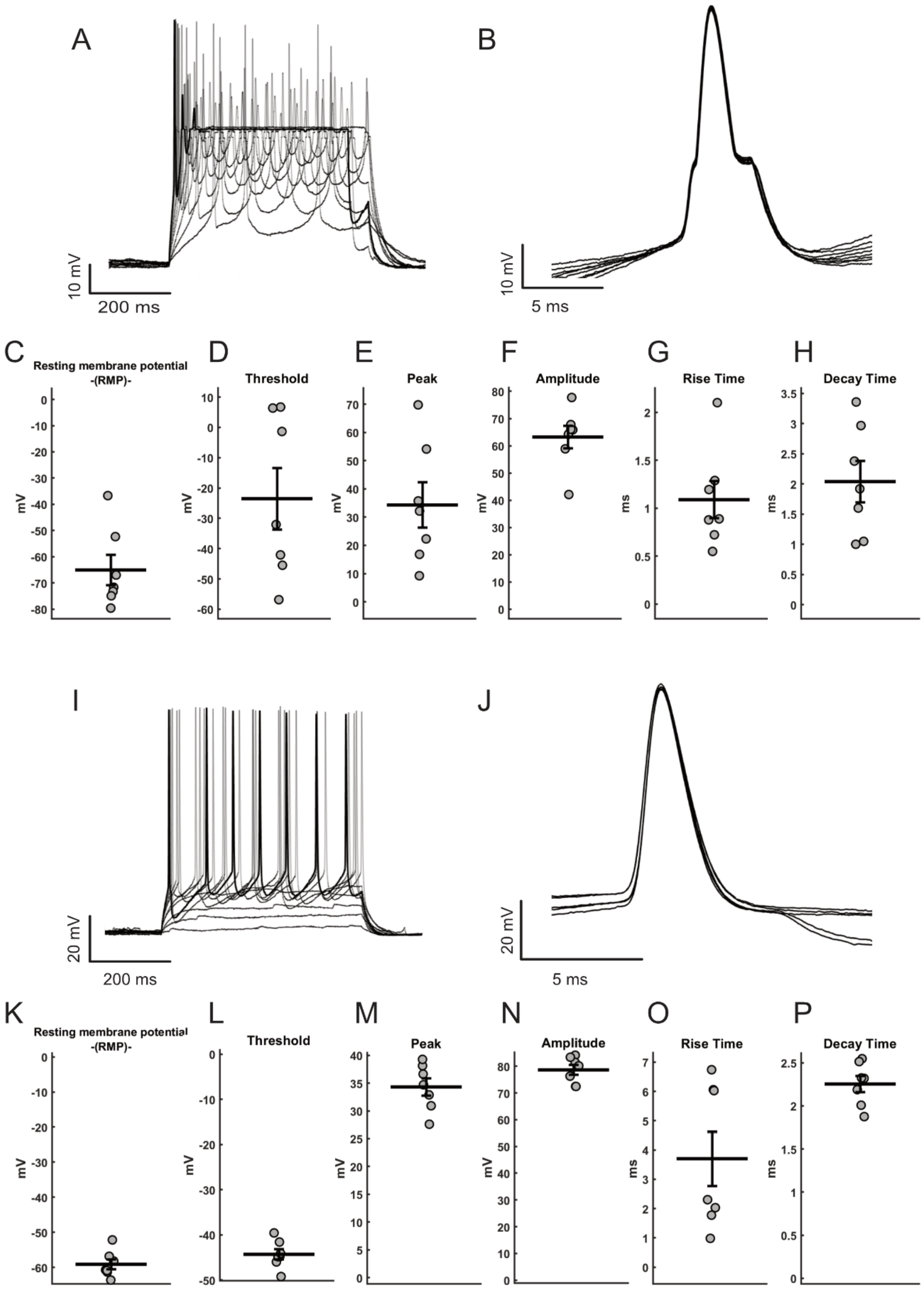


**Figure S5. Functional characterization of primary cortical neurons at timepoints of 14div (A-H) and 21div (I-P). (A, I)** Cell depolarization and action potential resulting from current injection (A: overlaid incremental depolarization and appearance of an action potential in a single cell across 100 pA current steps; I: overlaid incremental depolarization and appearance of an action potential in a single cell across 20 pA current steps). **(B,**

**J)** Overlaid action potentials (APs) from all the events recorded during a single measurement. **(C, K)** Resting membrane potential for 14div and 21div, respectively. **(D, L)** AP threshold. **(E, M)** Peak potential of AP. **(F, N)** Amplitude of AP (from threshold to peak). **(G, O)** Rise time of AP (10%-90% of amplitude). **(H, P)** Decay time of AP (90%-10% of amplitude). In graphs C-H and K-P the thick black line represents the average value over the population (14div, n= 7; 21div, n=7) while the grey dots represent the average value of the individual cells.

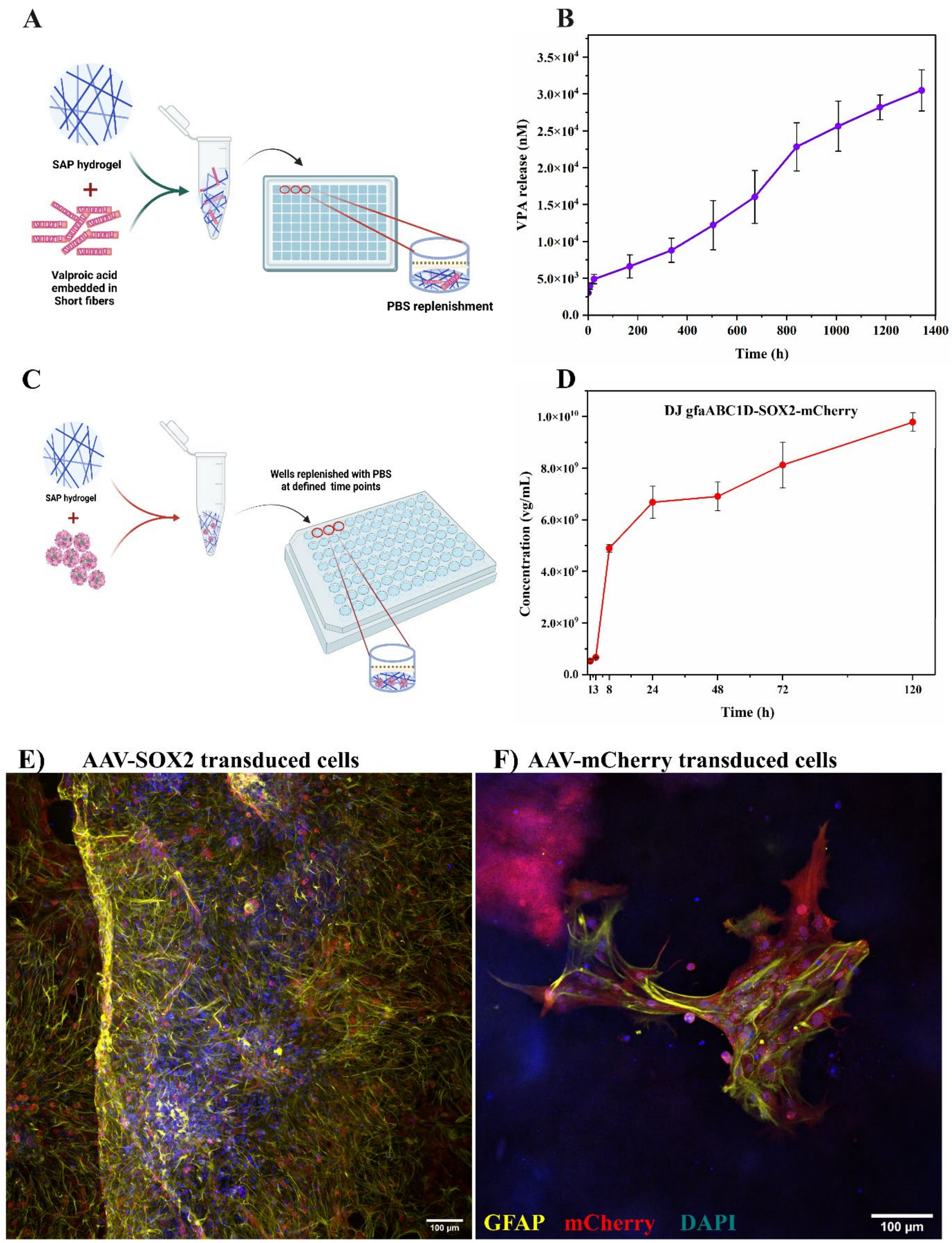


**Figure S6. Hydrogel encapsulated VPA/ and AAV-SOX2 prolonged payload release. A, C)** Schematic diagram of *in vitro* study design, **B, D)** Quantification of VPA and AAV-SOX2 release from over 8 weeks/ and 5 days, respectively. Representative images of **(E)** AAV-SOX2 and **(F)** empty vector (AAV-mCherry) transduced cells. Scale bar= 100 µm. Data represent mean ± SEM. n=3/time point

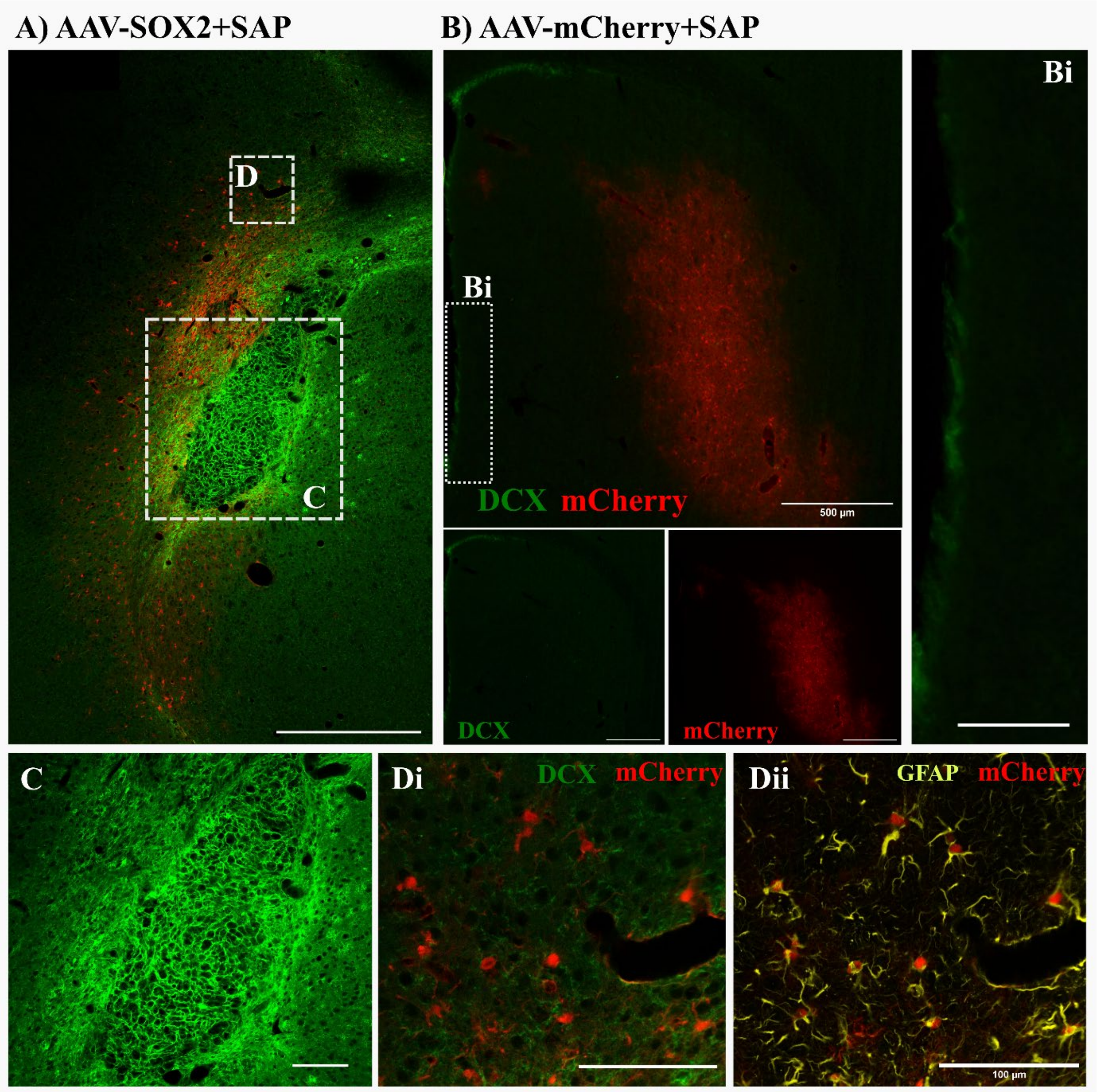


**Figure S7. SOX2 ectopic expression induces doublecortin (DCX+) neuroblast cells.** Representative images at the **A)** AAV-SOX2 + SAP and **B)** AAV-mCherry + SAP injection site illustrating DCX+ cells (green) and mCherry transduced cells (red), Scale bar= 500 µm. **C)** high magnification images of the core with DCX+ cells, **D)** High magnification image of the area slightly away from the injury site further confirming the constructed AAV preferentially transduce astrocytes. Scale bar (Bi, C, Di, Dii) = 100 µm.

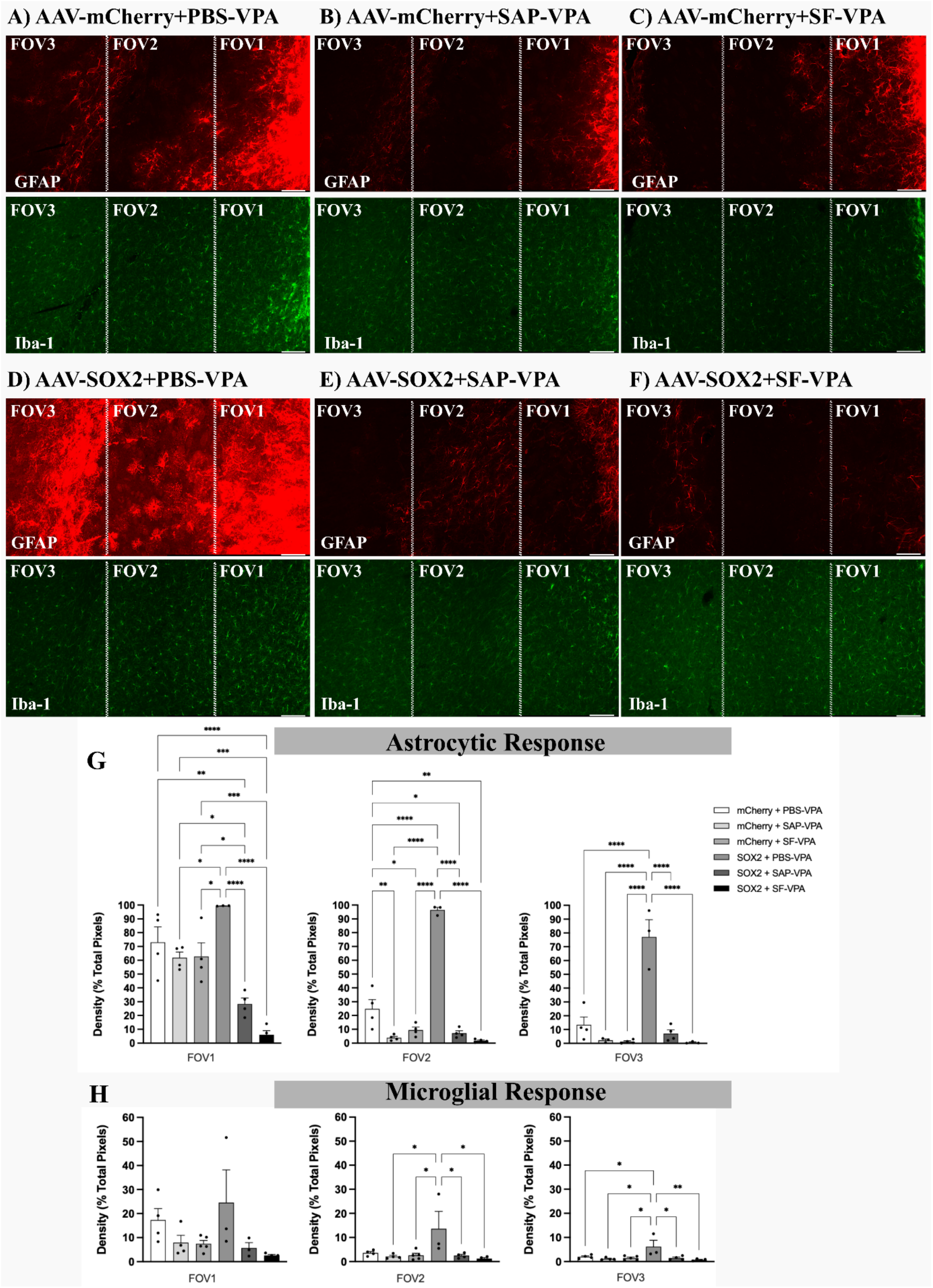


**Figure S8. The sustained release of AAVs through biomaterial attenuates the glial response in the mouse brain.** The different field of view (FOV) of representative coronal sections of different groups shows significant alleviation of inflammatory response and glial scar in AAV-SOX2 + SF-VPA treated groups. synergistic effect of AAV-SOX2 and composite biomaterial reduced glial scars and microglial inflammatory responses. **A-F)** Representative fluorescent images of the coronal sections of different groups stained for astrocytes (GFAP, red)

and microglia (Iba-1, green). Quantified measurements of the density of **(G)** astrocytes and **(H)** microglia. *$p < 0.05$, ****$p < 0.0001$. Data are represented as the mean ± SEM (n = 4−5 per group). Scale bar= 100 μm
